# Nonmuscle myosin IIA dynamically guides regulatory light chain phosphorylation and assembly of nonmuscle myosin IIB

**DOI:** 10.1101/2021.12.20.473479

**Authors:** Kai Weißenbruch, Magdalena Fladung, Justin Grewe, Laurent Baulesch, Ulrich S. Schwarz, Martin Bastmeyer

**Affiliations:** Zoological Institute, Karlsruhe Institute of Technology (KIT), 76131 Karlsruhe, Germany; Institute of Functional Interfaces (IFG), Karlsruhe Institute of Technology (KIT), 76131 Karlsruhe, Germany; Institute for Theoretical Physics, University of Heidelberg, Philosophenweg 19, 69120 Heidelberg, Germany; BioQuant-Center for Quantitative Biology, University of Heidelberg, Im Neuenheimer Feld 267, 69120 Heidelberg, Germany; Institute for Biological and Chemical Systems - Biological Information Processing (IBCS-BIP), Karlsruhe Institute of Technology (KIT), 76344 Karlsruhe, Germany

**Keywords:** Mechanobiology, cellular force generation, mechanosensing, actomyosin, nonmuscle myosin II, minifilaments, isoforms

## Abstract

Nonmuscle myosin II minifilaments have emerged as central elements for force generation and mechanosensing by mammalian cells. Each minifilament can have a different composition and activity due to the existence of the three nonmuscle myosin II isoforms A, B and C and their respective phosphorylation pattern. We have used CRISPR/Cas9-based knockout cells, quantitative image analysis and mathematical modelling to dissect the dynamic processes that control the formation and activity of heterotypic minifilaments and found a strong asymmetry between isoforms A and B. Loss of NM IIA completely abrogates regulatory light chain phosphorylation and reduces the level of assembled NM IIB. Activated NM IIB preferentially co-assembles into pre-formed NM IIA minifilaments and stabilizes the filament in a force-dependent mechanism. NM IIC is only weakly coupled to these processes. We conclude that NM IIA and B play clearly defined complementary roles during assembly of functional minifilaments. NM IIA is responsible for the formation of nascent pioneer minifilaments. NM IIB incorporates into these and acts as a clutch that limits the force output to prevent excessive NM IIA activity. Together these two isoforms form a balanced system for regulated force generation.

## Introduction

The ability to generate intracellular forces is an essential prerequisite during development and homeostasis across the animal kingdom, and affects cellular morphodynamics from the supracellular to the subcellular scale (LeGoff and Lecuit, 2015; Sellers, 2000; Vogel and Sheetz, 2006; Yusko and Asbury, 2014). In analogy to the force generating units in muscle syncytia, nonmuscle cells express nonmuscle myosin II (NM II) motor proteins, which assemble into bipolar arrangements, so-called minifilaments (in reference to their small size compared to their counterparts in muscle cells) (Dasbiswas et al., 2018). NM II minifilaments are the downstream effectors that ultimately generate the driving force for numerous cellular processes. In contrast to muscle sarcomeres, where force generation is mainly regulated via Ca^2+^/Calmodulin-dependent signaling (Kobayashi and Solaro, 2005; Szent-Gyorgyi, 1975; Wakabayashi, 2015), the spatiotemporal control of contractile forces in nonmuscle cells is facilitated via continuous assembly/disassembly cycles of NM II minifilaments that are mainly regulated by the small GTPase RhoA (Heissler and Sellers, 2016; Sandquist et al., 2006; Vicente-Manzanares and Horwitz, 2010).

Assembly of NM II molecules into its functional units is a hierarchical process (Garrido-Casado et al., 2021; Heissler and Sellers, 2016). The NM II holoenzyme is a hexamer consisting of two heavy chains (NMHC II) that form a homodimer, two regulatory light chains (RLCs), and two essential light chains (ELCs). In an inactive conformation (10S), the coiled-coil tails of NMHC IIs are folded at positions close to the skip residues interrupting the heptad repeat of the coiled-coil, and attached to the myosin heads, thus preventing their energy-burning cycling (Heissler and Manstein, 2013; Juanes-Garcia et al., 2016). Phosphorylation of the RLCs at Ser19 mediates the transition from the assembly-incompetent 10S to the assembly-competent 6S conformation (Billington et al., 2013). Although there is an additional phosphorylation at Thr18, which has an additive effect on the ATPase activity of the myosin heads, the phosphorylation at Ser19 is sufficient to facilitate the conformational unfolding (Garrido-Casado et al., 2021; Sellers and Heissler, 2019). In the final state, 28-30 individual NM II hexamers typically assemble in a bipolar minifilament with around 15 hexamers on each side and a typical size of 300 nm (Billington et al., 2013). The tail-tail interactions in the NMHC II homodimer are known to be mainly of electrostatic nature and depend on a characteristic charge pattern (+), which is present in the assembly-competence domains (ACD) (Dulyaninova and Bresnick, 2013) and allows for both, parallel and anti-parallel stacking (Kaufmann and Schwarz, 2020; Ricketson et al., 2010; Straussman et al., 2005). Because the block in the head region is now removed, the myosin II motors in the minifilaments are ready to cycle through the crossbridge cycle and thus to generate force.

To regulate the contractile output of the minifilaments more precisely, mammals express up to three different NM II isoforms simultaneously. The isoforms, commonly termed NM IIA, NM IIB, and NM IIC, have different heavy chains and differ with respect to their ATP-dependent motor activity, which is determined by their NMHC II head domains (Billington et al., 2013). Especially the ubiquitously expressed isoforms, NM IIA and NM IIB, possess complementary kinetics. While NM IIA possesses a higher ATPase activity, thus propelling actin filaments 3.5× faster than the other paralogs (Kovacs et al., 2003; Wang et al., 2000), NM IIB has an higher ADP affinity and can bear more load due to its higher duty ratio (Pato et al., 1996; Wang et al., 2003). The two isoforms also have different disassembly kinetics, which are mainly determined by their NMHC II tail domains and the non-helical tailpiece (NHT) (Dulyaninova and Bresnick, 2013; Dulyaninova et al., 2007; Dulyaninova et al., 2005; Heissler and Manstein, 2013). In particular, it has been shown that the subcellular localization patterns of NM IIA and NM IIB can be exchanged by swapping their tail domains (Juanes-Garcia et al., 2015; Sandquist and Means, 2008; Taneja et al., 2020).

Functionally, several studies have demonstrated that NM IIA acts as the first responder during force generation, while NM IIB supports this function by balancing and stabilizing the pre-initiated contractions on longer time scales (Taneja et al., 2020; Weissenbruch et al., 2021). Structurally, it was shown that the form follows the designated functions. With the advent of super-resolution microscopy, it became clear that the minifilaments in mammals are actually mixtures of the three different isoforms (Beach et al., 2014; Shutova et al., 2014) and that a gradient of NM IIB exists from the leading edge to the cell center of polarized cells (Beach et al., 2014), restricting the main activity to the more stable rear part of the cell body (Juanes-Garcia et al., 2015; Vicente-Manzanares et al., 2008; Vicente-Manzanares et al., 2011).

Despite these important advances, it is still not fully understood how activation, assembly and disassembly of the different isoforms are dynamically orchestrated to build up and maintain a polarized actomyosin cytoskeleton. Since all isoforms contain the same set of RLCs and ELCs, but vary with respect to their NMHC IIs (Heissler and Manstein, 2013), their activation would theoretically rely only on the quantitative ratio of the isoforms in the minifilament. However, independent studies suggest that the assembly of the actomyosin system is not only given by a random sequence of incorporation of new NM II hexamers, but follows a regulated sequence (Beach et al., 2014; Fenix et al., 2016; Shutova et al., 2017; Taneja et al., 2020; Weissenbruch et al., 2021). However, it is not clear yet how this dynamical sequence is established.

In an earlier study, we have introduced CRISPR/Cas9-generated NM II-KO cell lines and studied the influence of NM II isoforms on the cellular morphodynamics (Weissenbruch et al., 2021). Here we use these cell lines together with super-resolution microscopy, quantitative image analysis and mathematical modelling, to show that the amount of RLC phosphorylation is not equal to overall NM II activity, but that there is a strong asymmetry between NM IIA and NM IIB. The loss of NM IIA nearly completely abrogated the pRLC signal intensity, while the loss of NM IIB and NM IIC had no effect in this regard. Strikingly, only re-expressing NMHC IIA, but not overexpressing NMHC IIB, fully restored the RLC phosphorylation signals. Moreover, the assembly properties of NM IIB were influenced by the NM IIA-KO, while the NM IIB-KO had no effect on the assembly properties of NM IIA. Thus, the activation and assembly of initiating ‘pioneer’ hexamers is significantly higher for NM IIA than for NM IIB. NM IIB hexamers instead preferentially co-assemble into pre-formed NM IIA minifilaments. NM IIC minifilaments were only very sparsely associated with pRLC (Ser19) signals and NM IIA minifilaments, showing that this isoform might be part of a non-canonical molecular pathway. NM IIA and B in contrast perform together in a counterbalancing system that locally self-amplifies its activity via a positive feedback loop. Once assembled, NM IIB autonomously amplifies its dwell time in the minifilament, in analogy to a catch-bond mechanism, through force itself. This way, the load-bearing properties of the minifilaments prolong autonomously, assuring stable mechanotransduction in a noisy environment. At the same time, contractile overshoots from excessive NM IIA activity are prevented, here demonstrated by reinforcing the NM IIA activity in the heterotypic minifilaments via preventing the phosphorylation-induced disassembly of NM IIA hexamers.

## Results

### NM IIA amplifies the phosphorylation of RLCs at Ser19

To investigate the mutual interference of the NM II isoforms during the spatiotemporal assembly of the actomyosin system, we used U2OS cells, which express all three NMHC II isoforms simultaneously and are a widely used model system for the study of focal adhesions (FAs) and actin stress fibers (SFs) (Burnette et al., 2014; Hotulainen and Lappalainen, 2006; Jiu et al., 2019; Lee et al., 2018; Tojkander et al., 2015; Tojkander et al., 2011). Lately, we have generated stable knock out cell lines from these cells, which are deficient for one of the NMHC II paralogs, each (Weissenbruch et al., 2021). This collection allowed us to systematically investigate the influence of all three isoforms from the same cellular background.

The first key step during the assembly of the actomyosin system is the conformational extension of NM II hexamers via phosphorylation of the NMHC II associated RLCs at Ser19. Thus, we started by investigating the local and global correlation between Ser19 pRLCs and the various NMHC II isoforms in our cell lines in more detail. Using super-resolution AiryScan microscopy, we first visualized Ser19 pRLCs together with NMHC IIA, B, or C in WT cells, and compared their spatial correlation in bipolar minifilaments, as schematically depicted (Figure 1A). As expected, the pRLC signals strictly co-localized with the neck regions of NMHC IIA and NMHC IIB signals in the known bipolar fashion (Figure 1B), with NMHC IIB signals being especially enriched in the cell posterior, as shown previously (Beach et al., 2014; Kolega, 1998; Sandquist and Means, 2008; Shutova et al., 2012; Shutova et al., 2014) (Supplement figure 1). In contrast, NMHC IIC signals showed a remarkably low co-localization with pRLC signals in bipolar minifilaments. Although we found an overall cellular co-localization along the SFs, high resolution detail images showed that pRLC and single NMHC IIC clusters only occasionally resembled a bipolar arrangement (Figure 1B and Supplement figure 1). To confirm this finding, we compared the same staining pattern in cells, where higher NMHC IIC levels are present. We tested A431, A549, and HCT-116 cells, and observed the same outcome: despite an intracellular co-localization along SFs, bipolar arrangements of pRLC and NMHC IIC signals were only rarely observed (Supplement figure 2). These results indicate that NM IIC minifilaments somewhat differ from the classical contractile motors NM IIA and NM IIB, as also suggested by the variation in hexamer number and bare zone length (Billington et al., 2013) or its unusual function during traction force generation and tensional homeostasis (Weissenbruch et al., 2021). More importantly, however, it shows that the amount of pRLC signals cannot be directly translated to total NM II activity.

**Figure 1:**
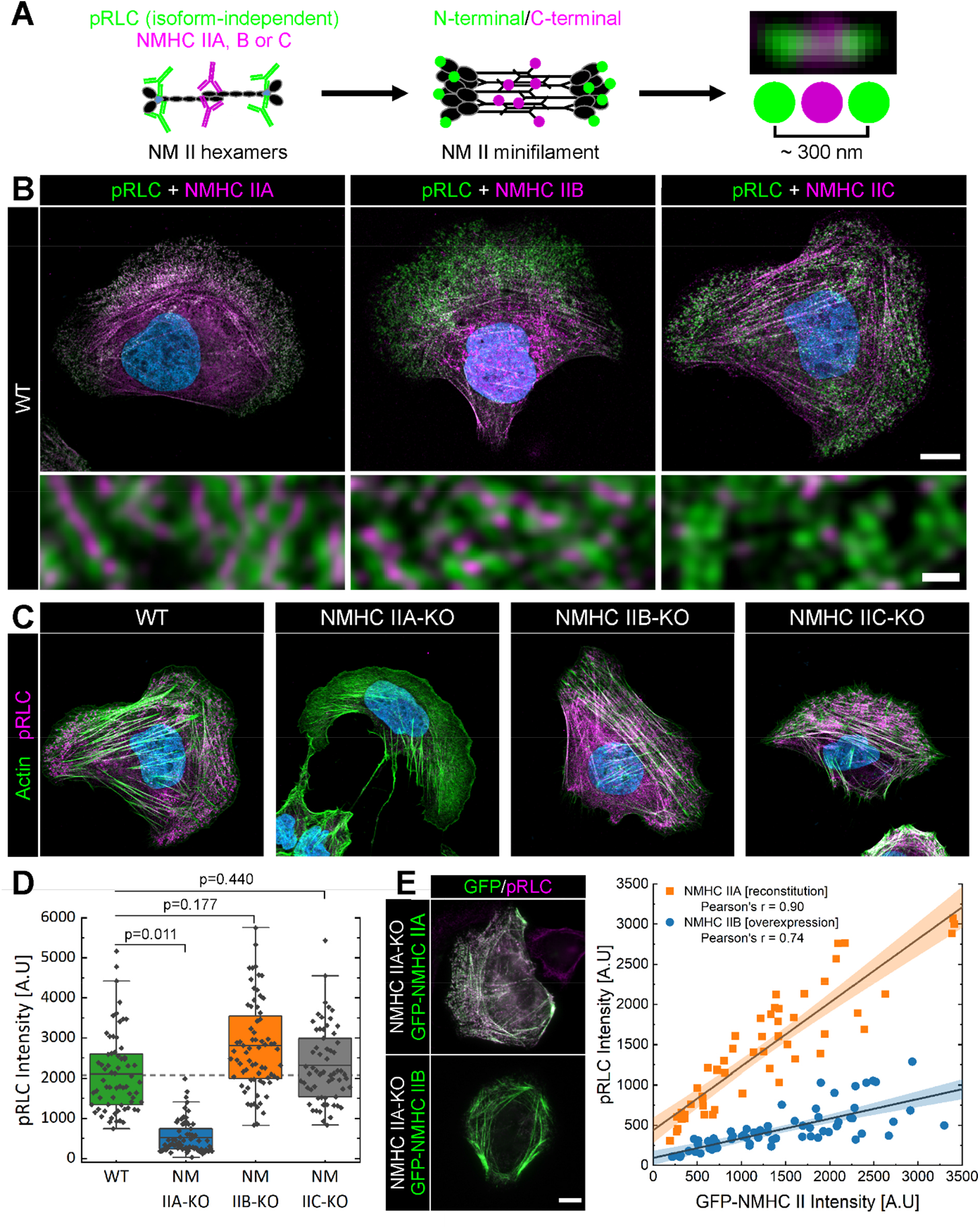
Expression of NMHC IIA is correlated with the phosphorylation of RLCs at Ser19. **(A)** Cartoon depicting the labeling strategy for NM II minifilaments. Antibodies against the Ser19 pRLCs theoretically label the head regions of all active minifilaments, while isoform-specific antibodies label the C-terminal region. **(B)** Fluorescence micrographs reveal a close spatial correlation for Ser19 pRLCs and NMHC IIA or NMHC IIB, while this is not the case for NMHC IIC. **(C)** Loss of NMHC IIA, but not NMHC IIB or NMHC IIC, abrogates the Ser19 pRLC level. **(D)** Ser19 pRLC signal intensity was quantified by measuring mean fluorescence intensity from line scans along segmented SFs. Values from three independent experiments (N=3) and n=30 cells of each cell line with up to three independent line scans per cell were plotted. **(E)** Reconstitution of NMHC IIA leads to a linear increase of Ser19 pRLC signal intensity (Pearson’s r = 0.90). Overexpression of GFP-NMHC IIB in NMHC IIA-KO cells also correlates with pRLC signal intensity (Pearson’s r = 0.74) but only shows a modest increase. Plots were derived from the following data sets: NMHC IIA = 32 line scans from 22 cells; NMHC IIB = 66 line scans from 32 cells. Scale bars represent 10 μm in overviews and 0.5 μm in insets of (B).

To quantitatively measure the impact of the different NM II paralogs on the global activation of the actomyosin system, we stained Ser19 pRLCs in the respective NMHC II-KO cell lines and quantified the signal intensities along segmented SFs (Figure 1C). Strikingly, the pRLC staining was almost completely absent in NMHC IIA-KO cells, although NMHC IIB and NMHC IIC were still expressed (Figure 1C&D). In contrast, it was not at all reduced in NMHC IIB-KO or NMHC IIC-KO cells and we even measured a slight increase in mean pRLC signal intensity for NMHC IIB-KO cells (Figure 1C&D). We then also stained for pan RLCs in NMHC IIA-KO cells and again found only a low fraction of RLC molecules assembled in the remaining minifilaments (Supplement figure 3A). Moreover, we also tested NMHC IIA-deficient COS-7 cells, which are known to express higher levels of NMHC IIB and NMHC IIC (Bao et al., 2005), and observed a comparable low pRLC signal (Supplement figure 3B).

Surprised by the strong independence of RLC phosphorylation from the concentration of NMHC II, we further tested the qualitative influence of the closely associated NMHC II isoforms A and B on the pRLC status. When we reconstituted NM IIA in NMHC IIA-KO cells by expressing GFP-tagged NMHC IIA, we observed a positive linear correlation between the pRLC signal intensity and NMHC IIA expression ratio (Figure 1E). In contrast, overexpression of GFP-tagged NMHC IIB in NMHC IIA-KO cells resulted in a significantly lower pRLC signal amplification, although NMHC IIB was expressed under the same constitutively active promotor, reaching NM IIB expression levels comparable to NM IIA (Figure 1E). No correlation between NMHC IIA expression ratio and pRLC intensity was observed, when we overexpressed NMHC IIA in NMHC IIB-KO cells (Supplement figure 4), indicating that the actomyosin system might be already in a saturated/fully activated status due to the presence of endogenous NM IIA.

Together, these results reveal that the different NM II isoforms have very different propensities to be associated with Ser19 pRLCs. This implies that pRLC as a marker only insufficiently reflects the complexity of the processes that guide the spatiotemporal assembly of the different NM II paralogs.

### NM IIA influences the assembly properties of NM IIB

The second step after the activation of the individual hexamers is their assembly in homotypic and/or heterotypic minifilaments. This raises the question how the different isoforms can be integrated into one system in a functional manner, if it relies on the same activation signal. Several groups reported that NM IIA and NM IIB co-assemble in heterotypic minifilaments (Beach et al., 2014; Shutova et al., 2014). Additionally, Beach and colleagues described the presence of heterotypic NM IIA/C minifilaments (Beach et al., 2014). However, in our case, this combination did by far not occur as regular as the combination NM IIA/B. Due to the low co-localization between pRLC and NMHC IIC, we found only very few bipolar arrangements that resembled heterotypic NM IIA/C minifilaments (Supplement figure 5A&B). We therefore focused in the following on the paralogs NM IIA and NM IIB, which are considered the main drivers for cellular contractility.

The loss of NM IIA not only abrogated the pRLC, level but also had a strong impact on the distribution of the remaining NM IIB minifilaments in the cell body. Compared to the gradual distribution along the cell axis in WT cells, the NM IIB minifilaments were now strikingly reduced in numbers and the remaining minifilaments densely clustered along the few remaining actin SFs (Figure 2A). The peak intensities of these single NMHC IIB assemblies were, however, up to 10-times higher in NMHC IIA-KO cells compared to WT cells (Figure 2B). We then measured the mean intensities of NM IIB minifilaments along segmented SFs in the cell periphery of WT and NMHC IIA-KO cells and found a 1.48-fold increase in overall NMHC IIB signal intensity in the absence of NMHC IIA (Figure 2C). When we compared the mean NMHC IIB intensities in the cell center, where the highest signal intensity for WT NM IIB minifilaments were observed, we still found a 1.3-fold increase (Figure 2C). Although we observed large spreads of intensities along the distinct SFs, these results indicate that more NM IIB hexamers clustered in a single minifilament, independent of the subcellular position.

**Figure 2:**
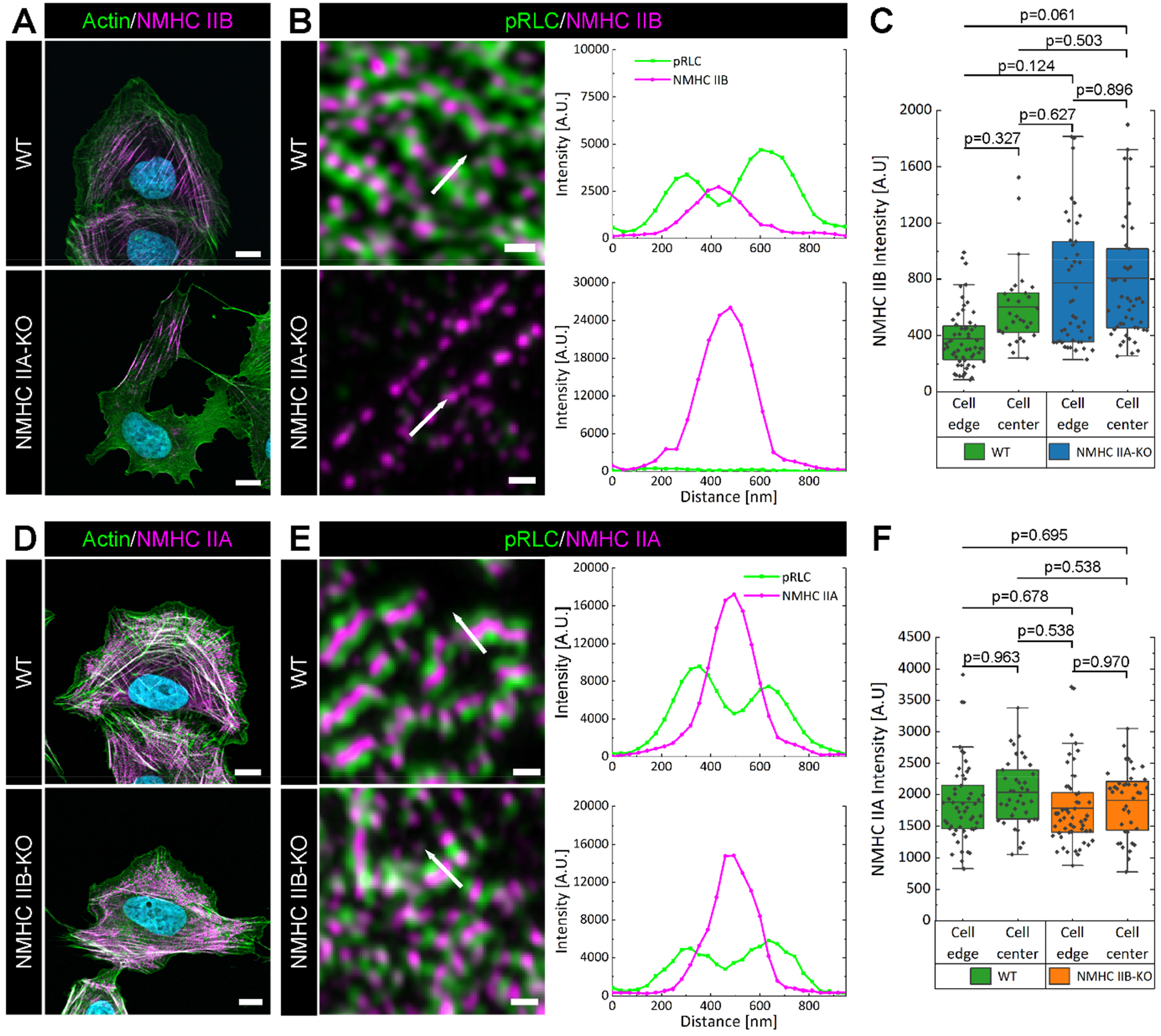
Loss of NM IIA decreases NM IIB density in minifilaments. **(A)** The localization of NM IIB is strongly affected by the loss of NM IIA. Few NM IIB minifilaments remain and their localization is restricted to small areas with densely compacted minifilament arrays. **(B)** High magnification images show single NM IIB minifilaments with up to 10-fold higher intensity values in the absence of NM IIA (compare line scans along the regions depicted by arrows). **(C)** NMHC IIB mean signal intensities were quantified by measuring fluorescence intensities from line scans along segmented SFs. **(D)** The localization of NM IIA is not affected by the loss of NM IIB. **(E)** Single NM IIA minifilaments contain comparable amount of NM IIA hexamers in the presence or absence of NM IIB, as indicated by the line scans in the depicted regions. **(F)** NMHC IIA mean signal intensities were quantified by measuring fluorescence intensities from line scans along segmented SFs. Values for (C) and (F) are derived from 90 line scans and 30 cells of each cell line. Scale bars represent 10 μm in (A) and (D), and 0.5 μm in (B) and (E).

In contrast, the assembly rate of NM IIA was not different in the presence or absence of NM IIB. Neither the intracellular distribution (Figure 2D), nor the intensity of single NM IIA clusters (Figure 2E), or the mean NM IIA intensity (Figure 2F) differed between WT and NM IIB-KO cells. Since NM IIA and NM IIB minifilaments both contain ~28-30 hexamers in *in vitro* assemblies (Billington et al., 2013; Niederman and Pollard, 1975), the lower number of NM IIB hexamers in minifilaments of WT cells suggests that binding sites might be occupied by NM IIA, rendering heterotypic NM IIA/B minifilaments the basic contractile unit in mammalian cells. Vice versa, however, the loss of NM IIB did not change the assembly properties of NM IIA, suggesting that NM IIA acts ‘upstream’ of NM IIB during the assembly of heterotypic minifilaments.

### NM IIA guides the frequency and spatial distribution of NM IIB in the cell body

We next checked how the subcellular distribution of heterotypic NM IIA/B minifilaments is influenced by the reconstitution of NM IIA. It was suggested that pre-ordered NM IIA filament stacks might serve as a template for the co-assembly of NM IIB hexamers (Fenix et al., 2016; Shutova et al., 2017). To investigate this hypothesis in our system, we expressed C-terminally tagged NMHC IIA-mApple in NMHC IIA-KO cells and stained for endogenous NM IIB. Since both fluorophores are located in the C-terminal regions of the target isoform, overlapping signal spots visualize heterotypic minifilaments with high precision, when imaged with super-resolution AiryScan or SIM microscopy.

Reconstitution of NM IIA restored the frequency and distribution of NM IIB in the cell body (Figure 3A). In cells that adopted a polarized phenotype, NM IIA was homogenously distributed throughout the cell body, while NM IIB was enriched at the cell center (Beach et al., 2014; Kolega, 1998; Shutova et al., 2012; Shutova et al., 2014). Comparing the signal intensities of NM IIA and NM IIB in single clusters showed that the leading edge was decorated with nascent NM IIA minifilaments, which contained only a very low amount of NM IIB hexamers, if they were detectable at all (Figure 3B). With progressing distance from the leading edge, more NM IIB hexamers co-assembled into the NM IIA pioneer minifilaments, until a balanced ratio of NM IIA and NM IIB was reached in the center of the cell and along the trailing edge (Figure 3B). While the ratio of NM IIA and B changed with regard to the subcellular position in polarized cells, it remained constant in non-polarized cells (Figure 3C). This suggests that the gradual enrichment of NM IIB in the cell center of polarized cells might be facilitated via transport of bound NM IIB hexamers with the retrograde actin flow. The induction of heterotypic NM IIA/B minifilaments, however, occurs independent of the subcellular position, showing that the initiation is an intrinsic feature of NM IIA, as previously suggested (Shutova et al., 2014).

**Figure 3:**
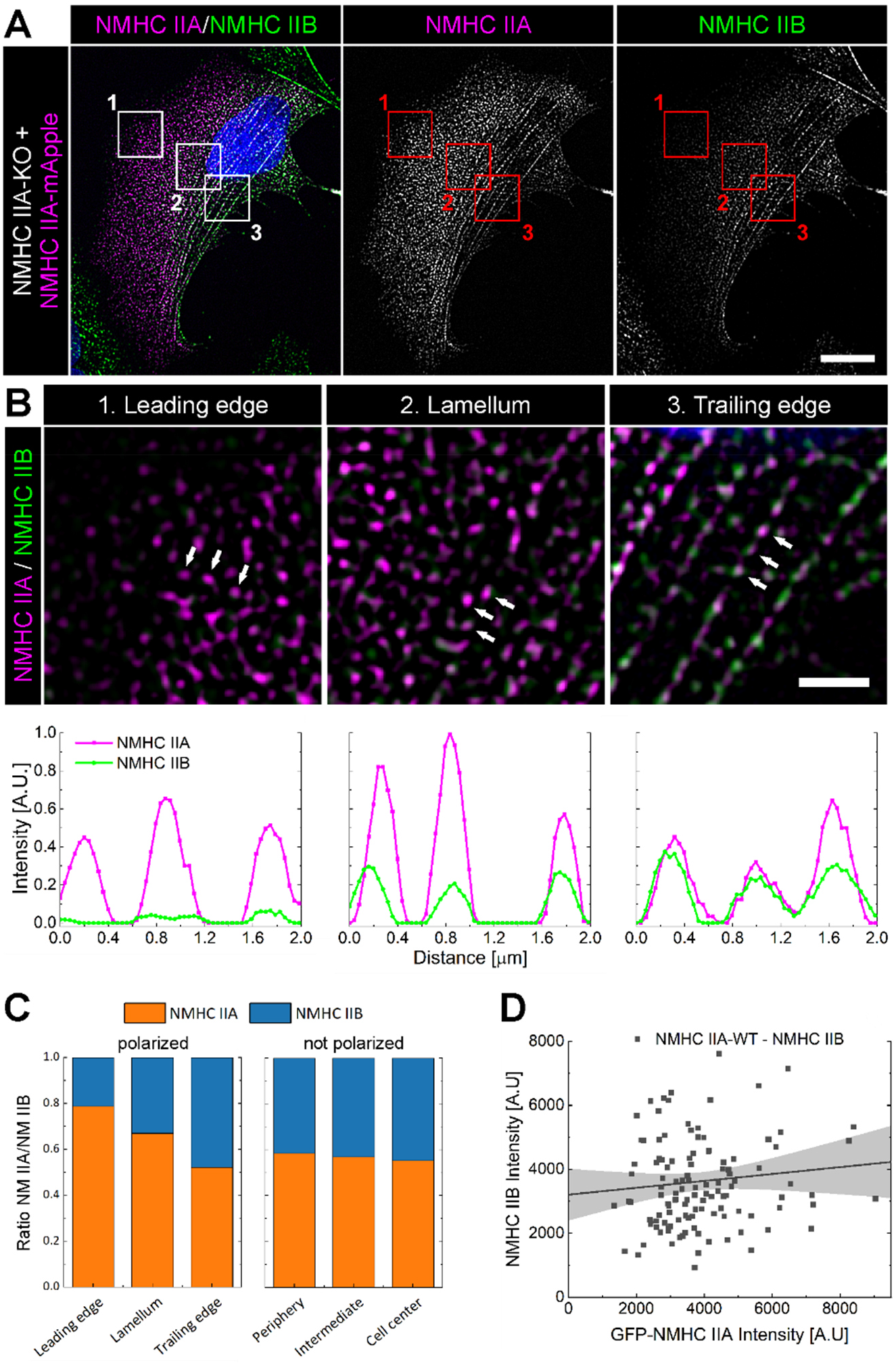
NM IIA templates for NM IIB and guides its distribution in the cell body. **(A)** NMHC IIA-KO cells were reconstituted with C-terminal tagged NMHC IIA-mApple (magenta) and endogenous NM IIB was labeled by an antibody recognizing the C-terminal tailpiece of NMHC IIB (green). Co-localization of NM IIA and NM IIB clusters were analyzed along the cell axis of polarized or non-polarized cells. Representative measurements were taken in three different spots as indicated by the red squares corresponding to 1-3. **(B)** In polarized cells, nascent NM IIA clusters were observed along the leading edge (1) and co-localization with NM IIB clusters was very low. With increasing distance from the leading edge, NM IIB co-assembles into NM IIA minifilaments (2) until a balanced ratio of NM IIA and NM IIB in heterotypic minifilaments is achieved in the cell center (3). Individual line scans depict co-localization of NM IIA and NM IIB in the three clusters, corresponding to the white arrows. **(C)** While the ratio of NM IIA and NM IIB changed with regard to the centripetal actin flow in polarized cells, it remained constant in non-polarized cells. **(D)** Plotting the intensities of NMHC IIA-mApple and NMHC IIB clusters shows that no quantitative correlation exists between the assembly ratio of both paralogs. For (C), line scans along SFs were derived from 16 individual polarized cells out of three independent experiments (N=3). For (D), plots were derived from 120 line scans of 30 cells. Scale bars represent 10 μm in (A) and 0.5 μm in (B).

Finally, we checked whether a quantitative correlation between the two paralogs might exist. However, in contrast to the spatial correlation, we found no correlation between the quantities of assembled NM IIA and NM IIB molecules in the heterotypic minifilaments (Figure 3D). Thus, although NM IIA guides NM IIB assembly, the dwell time and disassembly dynamics of the NM II hexamers are regulated in a paralog-specific manner.

### Force regulates the exchange dynamics of NM IIB hexamers in minifilaments

During the mature phase of the minifilament lifetime cycle, differential assembly/disassembly rates have to be established to maintain the self-sorting of the heterotypic minifilaments in a fully-polarized actomyosin cytoskeleton. We therefore asked next, how the different exchange dynamics of the paralogs are conducted, after the initiation and assembly of the minifilament is completed. Our previously described mathematical model for minifilament dynamics suggests that the crossbridge cycling rates of the different NM II isoforms directly translate into different minifilament stability (Grewe and Schwarz, 2020a). Given the known crossbridge-cycle rates, the assembly/disassembly of NM IIA should not be impeded much by force, corresponding to a slip-bond nature, while the disassembly of NM IIB should be impeded by force, corresponding to a catch-bond behavior (Grewe and Schwarz, 2020a; Grewe and Schwarz, 2020b). Thus, in analogy to the catch-bond mechanisms described for a subset of the integrins in FAs, force itself might induce the autonomous assembly of the actomyosin system.

To test this idea, we investigated how force influences the exchange dynamics of the individual hexamers by performing fluorescence recovery after photobleaching (FRAP) studies on both paralogs. We again reconstituted GFP-NMHC IIA or GFP-NMHC IIB in the respective NMHC II-KO cell line (Figure 4A&B and Movies 1&2) and measured their dynamics in the absence or presence of photostable para-aminoblebbistatin (Varkuti et al., 2016) (Figure 4C). When comparing the recovery times and mobile fractions in the absence of blebbistatin, we found that the exchange rate of NM IIA is much faster compared to NM IIB (Figure 4D&E). While NM IIA shows a mobile fraction of 63 ± 29% with an exchange timescale of 69 ± 53 s, NM IIB possesses a lower mobile fraction of 47 ± 28% with a higher exchange timescale of 230 ± 140 s. These results are in line with previously published results from other groups that measured FRAP dynamics by overexpressing NM IIA or NM IIB in different cell lines (Sandquist, 2008; Shutova et al., 2017; Vicente-Manzanares et al., 2008; Vicente-Manzanares et al., 2007). In the presence of blebbistatin, however, we find that NM IIA and NM IIB possess the same recovery dynamics for both, mobile fraction and recovery time (Figure 4C-E). For NM IIA in the presence of blebbistatin, we find a mobile fraction of 55 ± 34% with recovery timescales of 52 ± 30 s, while for NM IIB, a mobile fraction of 64 ± 33% with recovery timescales of 62 ± 44 s were observed. Since recovery timescale and mobile fraction are not statistically independent variables as they arise from the same fit, we compared the joint distribution of both observables by a two-dimensional version of the Kolmogorov-Smirnoff test, the Peacock test (Fasano and Franceschini, 1986; Peacock, 1983). Comparing our different conditions revealed significant differences only between NM IIB in the absence of para-aminoblebbistatin and all other experimental situation (Figure 4C-E and Table 1). Thus, blebbistatin had a strong impact on NM IIB but not on NM IIA.

**Figure 4:**
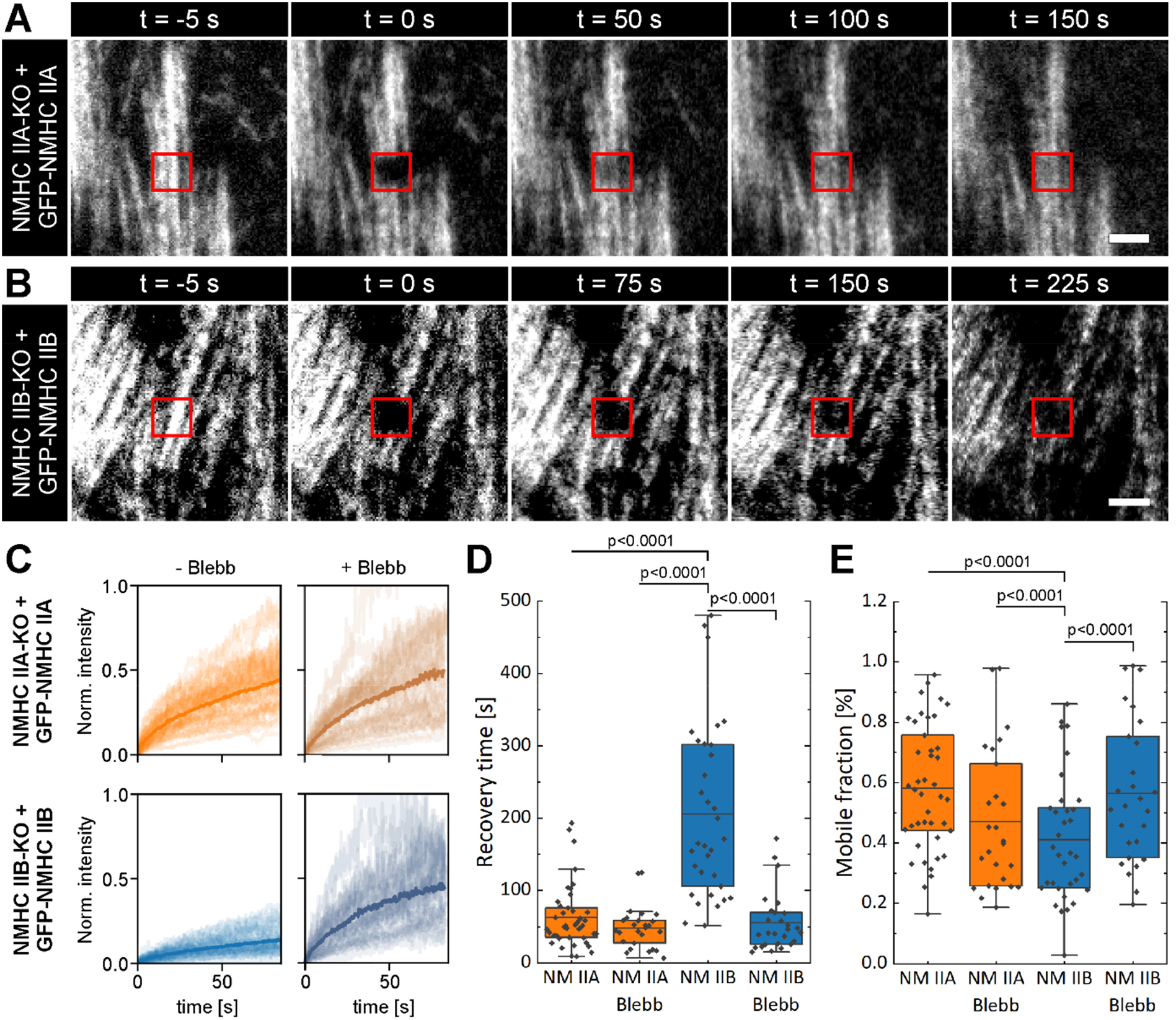
The exchange rate of NM IIB but not NM IIA is influenced by blebbistatin. **(A**) NM IIA or **(B)** NM IIB protein function was restored by reconstituting the GFP-tagged NMHC II isoform in the respective NMHC II-KO cell line. **(C)** Single recovery curves (thin lines) and average (thick line) for GFP-NMHC IIA or GFP-NMHC IIB in the absence or presence of photostable para-aminoblebbistatin (Blebb). **(D)** Boxplots showing the recovery time and **(E)** mobile fraction of GFP-NMHC IIA or GFP-NMHC IIB in the absence or presence of para-aminoblebbistatin. The recovery time of NM IIA is significantly faster compared to NM IIB and the mobile fraction is lower in the case of NM IIB. Treatment with blebbistatin abolishes these differences by significantly reducing the recovery time and increasing the mobile fraction of NM IIB but not NM IIA. Recovery times and mobile fractions were calculated from n_IIA_ = 45; n_IIA-Bleb_ = 30; n_IIB_ = 37 and n_IIB-Bleb_ = 30 individual traces and three independent experiments (N=3). Only p-values < 0.05 are shown and the complete evaluation is listed in table 1. Scale bar represents 10 μm for (A) and (B)

**Table 1:** p-values for FRAP measurements in Figure 4.

| Matrix 1 | Matrix 2 | p-value |
| --- | --- | --- |
| NM IIA-GFP | NM IIA-GFP + 50 μM Blebbistatin | 0.108495 |
| NM IIA-GFP | NM IIB-GFP | 5,10E+07 |
| NM IIA-GFP | NM IIB-GFP + 50 μM Blebbistatin | 0.333059 |
| NM IIA-GFP + 50 μM Blebbistatin | NM IIB-GFP | 1,47E+08 |
| NM IIA-GFP + 50 μM Blebbistatin | NM IIB-GFP + 50 μM Blebbistatin | 0.350563 |
| NM IIB-GFP | NM IIB-GFP + 50 μM Blebbistatin | 4,02E+09 |

Blebbistatin and presumably also its derivatives are known to target the tension generation of myosin II (Kovacs et al., 2004). The force-generating step of the crossbridge cycle, which is linked to phosphate release, is slowed down. As the differences in the FRAP experiments between NM IIA and NM IIB, which probe the assembly dynamics of myosin minifilaments, are leveled in the presence of blebbistatin, we interpret our result as evidence for an interdependence of the assembly of NM II minifilaments and their mechanochemical crossbridge cycle, resulting in a minifilaments that can tune its contractile output, i.e. its transition from slip- to catch-bond behavior in regard to the relative composition of NM IIA to NM IIB.

Aiming for a mechanistic understanding of this effect, we next used our NM II minifilament assembly model that couples the crossbridge cycle and the dynamic self-assembly to simulate the FRAP data (Grewe and Schwarz, 2020a; Grewe and Schwarz, 2020b). In brief, to model the association and dissociation dynamics, we start with a consensus architecture of the ~30 NM II hexamers that form a minifilament. Minifilaments are known to result from a very stable anti-parallel stagger that is complemented at the sides by parallel staggers (Figure 5A). In three dimensions, three such arrangements form a cylindrical structure that can be represented by an appropriate graph (Figure 5B). The presence or absence of fluorescent species can be simulated by using appropriate labels. The neighborhood relations of each NM II (lines in the graph) give rise to specific binding energies determined mainly by the electrostatic interactions of charged regions on the coiled-coils of the NM II hexamers (Kaufmann and Schwarz, 2020). All NM II hexamers in the assembly can interact with actin via the crossbridge cycle, thereby producing force. The presence of blebbistatin is reflected in a strong reduction of the rate at which the powerstroke occurs (rate k_12_ in Supplement figure S1). In our computer simulations, we assume that NM II hexamers cannot detach from the minifilament while they are part of the assembly (more details on the model and model parameters in the supplemental text).

**Figure 5:**
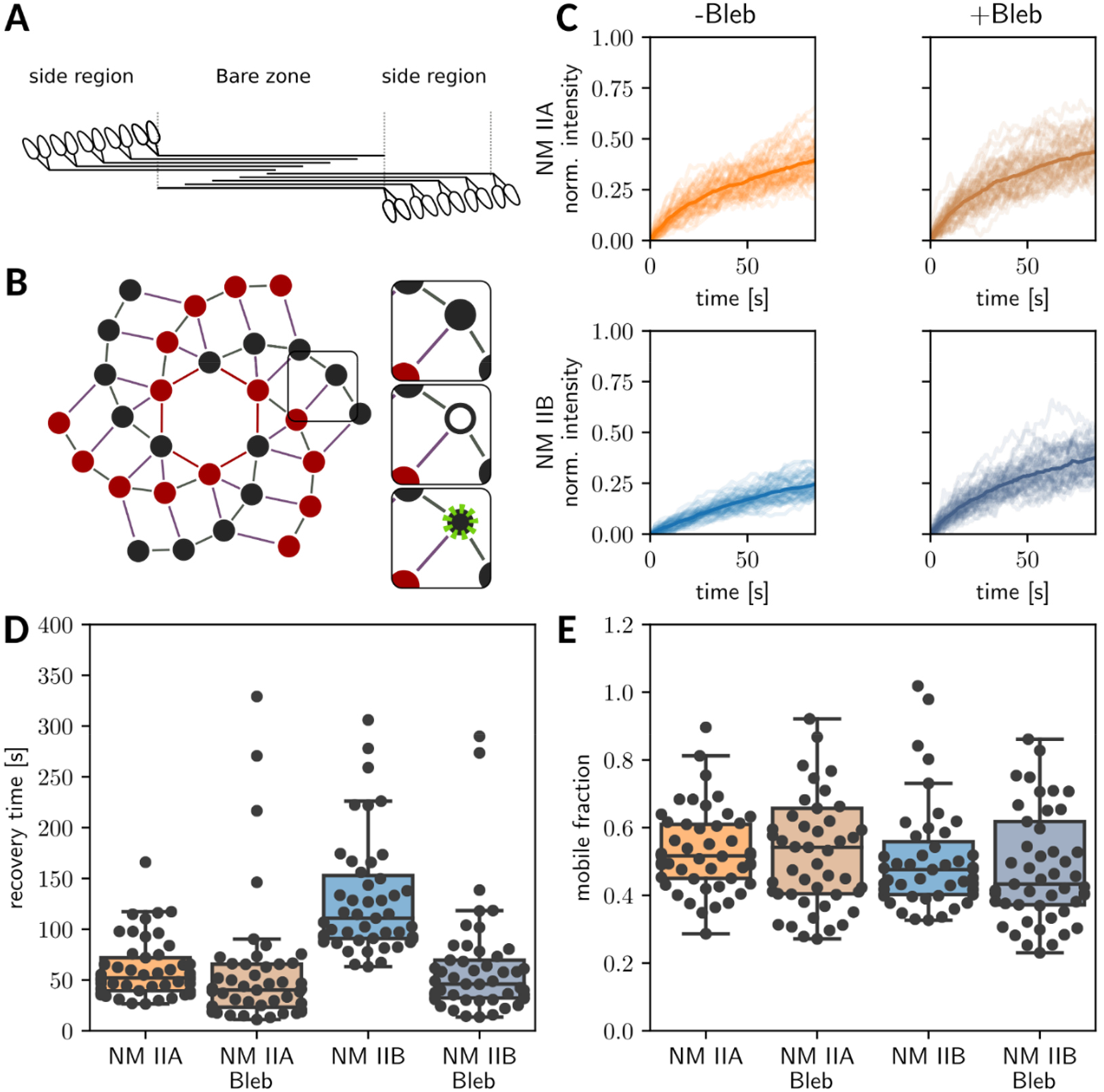
Stochastic computer simulations of the FRAP-experiments partially confirm the putative roles of the crossbridge cycling. **(A)** 2D slice through the consensus architecture for NM II minifilaments. **(B)** Graphical representation of the full 3D NM II minifilament. Each node represents one myosin II hexamer and each line represents a neighborship relation. The two different colors represent the two possible orientations. During FRAP, a bleached hexamer dissociates and a fluorescent hexamer enters. **(C)** Simulations of FRAP in the presence and absence of blebbistatin for mixed minifilaments with either NM IIA or NM IIB being fluorescently tagged. Blebbistatin is assumed to decrease the crossbridge cycle rate k_12_ (Supplement figure S1). **(D)** Recovery time and **(E)** mobile fraction predicted by the computer simulations.

We started by simulating NM IIA minifilaments in the absence of blebbistatin, which we used to calibrate the association rate to the experimental data. We obtained reasonable agreement with an association rate of *k*_*on*_ = 5 s^−1^, which we held constant in the following simulations. The simulated FRAP data are shown in Figure 5C. Simulating NM IIA in the presence of blebbistatin showed little change, consistent with the experiment. Simulating NM IIB with its slower detachment from the post-powerstroke state showed slower recovery dynamics that became comparable with the results for NM IIA. The timescales and mobile fractions are summarized in Figures 5D and E. We note that the NM IIB recovery times do not quantitatively match the experiments. Using an even slower detachment rate from the post powerstroke state gives better results, suggesting that the NM IIB post-powerstroke detachment may depend even more strongly on force than assumed in our current model. Overall, however, our model is able to qualitatively capture the effects that blebbistatin has on the FRAP dynamics. Thus, our data suggest that tension inhibits the exchange of NM IIB from the minifilament complex, causing NM IIB to move over longer periods of time with the retrograde actin flow and accumulate in the cell center.

### A counterbalancing system of NM IIA and NM IIB facilitates the force output

Once the NM IIA-induced assembly is initiated, the minifilaments start to generate force by crossbridge cycling. At the same time, NM IIB is incorporated into the minifilaments. If the amount of force exceeds a certain threshold, it activates the catch-bond behavior in heterotypic minifilaments by prolonging the duty ratio of NM IIB, thus autonomously stabilizing the contraction. NM IIB with its increased duty ratio simultaneously balances the tension and prevents overshoots, as they could be generated by excessive crossbridge cycling of NM IIA. Such a self-regulating circuit could maintain a homeostatic balance of tension, allowing cells to rapidly and stably transduce mechanical signals in a noisy mechanical environment. This view also suggests that NM IIB is automatically accumulated in the retrograde flow from the front to the back, simply by its longer dwell times.

To further test our interpretations, we tuned the crossbridge cycling ratio in the heterotypic complexes towards the kinetics of NM IIA by interfering with the assembly/disassembly kinetics. We restored NM IIA in NMHC IIA-KO cells, using three NMHC IIA mutants. NMHC IIA-ΔIQ2 lacks the binding sites for the RLCs, causing the NM IIA molecule to be constitutively in the assembly-competent conformation (Breckenridge et al., 2009). NMHC IIA-ΔNHT lacks the non-helical tailpiece of NMHC IIA, preventing its phosphorylation at S1943 (Breckenridge et al., 2009). In the mutant NMHC IIA-3xA, serine to alanine substitutions prevent phosphorylation of the C-terminal region on both prominent p-sites, S1943 and S1916 (Rai et al., 2017). Since phosphorylation of the NHT and coiled-coil region are believed to force the disassembly of the NM IIA hexamer from the minifilament complex (Breckenridge et al., 2009; Dulyaninova et al., 2007; Dulyaninova et al., 2005; Garrido-Casado et al., 2021), both mutants should prolong the dwell time of NM IIA and lead to an over-assembly of NM IIA in the minifilaments. As controls, we reconstituted NM IIA-KO cells with NMHC IIA-WT and an assembly-deficient NMHC IIA mutant, lacking the ACD domains (NMHC IIA-ΔACD). Thus, cells expressing the latter mutant should behave like untransfected NM IIA-KO cells.

All mutants, except for the assembly-incompetent NMHC IIA-ΔACD, resulted in the formation of heterotypic NM IIA/B minifilaments (Supplement figure 6A). NMHC IIA-ΔIQ2 expression did not change the pRLC level, although the distribution of NM IIB in the cell body was restored. Expression of both C-terminal phosphorylation-deficient mutants showed a positive linear correlation with pRLC signal intensities (Supplement figure 6B). However, compared to NMHC IIA-ΔIQ2, NMHC IIA-ΔNHT and NMHC IIA-3xA showed increased NMHC IIA signal ratios (Supplement figure 6C), suggesting an over-assembly of NM IIA in the heterotypic minifilaments.

To precisely quantify the influence of the NM IIA over-assembly on the mechanical phenotype, the cells were cultivated on cross-shaped micropatterns and their shapes were analyzed with our previously described dynamic tension-elasticity model (dTEM) (Weissenbruch et al., 2021). Briefly, the cell contour follows a sequence of inward-bent actin arcs with a circular shape that results from the interplay of two NM II-dependent tension regimes: the surface tension σ arises from the actin cortex and basically increases the curvature of the cell contour with increasing contractile strength, while the line tension λ arises in the actin arcs itself and decreases the curvature of the cell contour with increasing contractile strength. Balancing both results in a circular actin arc with the radius *R* = λ/σ (Laplace law). Since *R* is positively correlated with the spanning distance *d* between two adhesion sites (Bischofs et al., 2008), an elastic line tension λ(*d*) was implemented to explain the dependency on absolute distances. Importantly, we assigned NM IIA as the main generator of tensile forces, while NM IIB provides the elastic stability to the actin arcs (Weissenbruch et al., 2021). While the loss of NM IIA was accompanied with smaller arc radii due to low tension, the loss of NM IIB caused the formation of larger arc radii, which were not correlated to the spanning distance anymore.

We now have performed the same analysis for our NMHC IIA constructs. Reconstituting NM IIA protein function using NMHC IIA-WT revealed a positive *R*(*d*) correlation of 69 ± 6% (Figure 6A), which was nearly identical to what we observed for WT cells in our previous publications (compare black dotted line and solid red line) (Weissenbruch et al., 2021). Similar, expressing NMHC IIA-ΔACD does not alter *R*(*d*) correlation (71 ± 6%) and the value was comparable to untransfected NM IIA-KO cells in our previous work (compare black dotted line and solid purple line) (Supplement figure 7B). We then compared these values to the over-assembling mutants. Although NMHC IIA-ΔIQ2 accumulated in the central and rear part of polarized cells on homogenously coated substrates (Breckenridge et al., 2009) (compare Supplement figure 6A), it showed a comparable localization to NMHC IIA-WT on our micropattern, where the cells are in a steady-state. In accordance, also the *R*(*d*) correlation value was only slightly lower (61 ± 6%) compared to the WT control (Supplement figure 7C). However, when we tested the C-terminal phosphorylation mutants, we observed a significantly reduced *R*(*d*) correlation. This effect was observed when preventing phosphorylation at S1943 by expressing NMHC IIA-ΔNHT (53 ± 7%) (Figure 6B), and was even stronger when phosphorylation was prevented on both prominent p-sites, S1943 and S1916, by expressing NMHC IIA-3xA (43 ± 9%) (Figure 6C). Close inspection of the cellular phenotypes and the data revealed that the lower *R*(*d*) correlation results from the presence of almost straight arcs, which arise independent from the spanning distance, thus spreading out the data points. The same results were obtained in our previous publication, when NM IIB was depleted (Weissenbruch et al., 2021). This time, however, NM IIB is still expressed but the over-assembly of NM IIA nevertheless disrupts the *R*(*d*) correlation.

**Figure 6:**
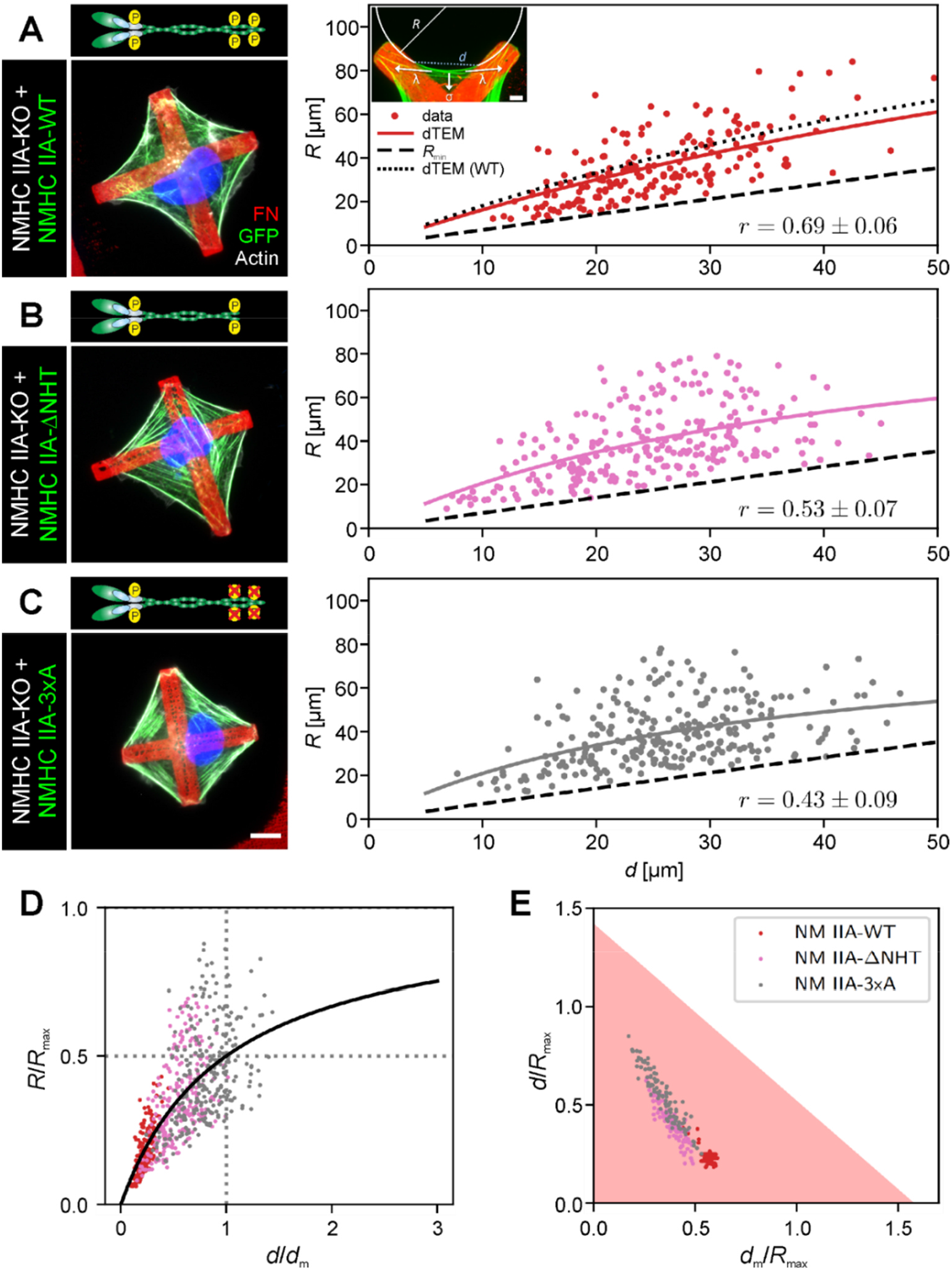
The ratio of NM IIA to NM IIB influences the force output of heterotypic minifilaments. Transfected cells were seeded on cross-shaped micropatterns and the *R*(*d*) correlation was analyzed (correlation coefficient r given at bottom right). Solid lines denote the bootstrapped mean fit of the dynamic tension-elasticity model (dTEM), black dashed lines denote the geometrically possible minimal radius. **(A)** Restoring NM IIA-WT protein function resulted in a similar *R*(*d*) correlation as for WT cells (compare black dotted line). **(B)** Expressing NMHC IIA-ΔNHT leads to a reduced *R*(*d*) correlation. **(C)** Expressing NMHC IIA-3xA leads to an even stronger reduction in *R*(*d*) correlation. **(D)** Rescaling the experimental values using the fit parameters shows that NMHC IIA-ΔNHT and NMHC IIA-3xA shifts the data points closer to the plateau regime of the dTEM master curve, while the data for NMHC IIA-WT lie in the linear regime. **(E)** The ratio *d*_*m*_/*R*_*max*_, which scales linearly with the ratio of SF friction and motor stall force, shows the same distribution with NMHC IIA-ΔNHT and NMHC IIA-3xA being distributed farest away from the minimal possible radius. Quantifications were derived from three independent experiments (N=3) with n_WT_ = 54; n_ΔNHT_ = 70 and n_3xA_ = 72 cells. Scale bar represents 10 μm in (A)-(C).

To further verify this observation, we plotted our results in the two-dimensional space of the dTEM. Fitting the equation

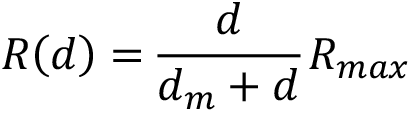

to the experimental data shown yield the parameters *R*_*max*_ and *d*_*m*_ for each NMHC IIA construct (Table 2). Rescaling the experimental values using the fit parameters shows that the data points of NMHC IIA-WT, NMHC IIA-ΔACD and NMHC IIA-ΔIQ2 are in the linear regime (Supplement figure 7D), while NMHC IIA-ΔNHT and NMHC IIA-3xA are shifted towards the plateau regime of the master curve (Figure 6D). While in the linear regime, the large *d*_*m*_ values correspond to high motor friction (known for NM IIB/large duty ratios), the plateau region is characterized by small *d*_*m*_ values that correspond to low motor friction (known for NM IIA/small duty ratios). Since NM IIB is required to elastically stabilize the correlation between arc radii *R* and spanning distance *d*, this suggests that the measured radii for NMHC IIA-ΔNHT and NMHC IIA-3xA are larger because *R*_*max*_ is realized by the enlarged dwell time of NMHC IIA molecules in the heterotypic minifilaments. Thus, the motor friction of the NMHC IIA molecules overpowers the actin flow out of the FAs, leading to the correlation breakdown and a tensile overshoot.

To further separate the different phenotypes, we plotted our data in the two-dimensional parameter space of (*d*_*m*_/*R*_*max*_, *d*/*R*_*max*_) (Figure 6E and Supplement Figure 7E). The shaded region denotes allowed values due to the central angle being smaller than 90°. Recently, we showed that the ratio *d*_*m*_/*R*_*max*_ increases with the relative amount of NM IIB, from NM IIB-KO over WT to NM IIA-KO cells. Strikingly, we observed a roughly similar sorting for our NMHC IIA variants. The ratio *d*_*m*_/*R*_*max*_ was lowest for NMHC IIA-3xA and NMHC IIA-ΔNHT (Figure 6E), increased for NMHC IIA-WT and NMHC IIA-ΔIQ2 and was highest for NMHC IIA-ΔACD (Supplement figure 7E). Thus, tuning the kinetics of NMHC IIA towards lower disassembly rates resembled mechanical features of SFs in NM IIB-KO cells, likely caused by an overshoot of NM IIA-derived tension, upstream of NM IIB.

**Table 2:** mean bootstrapped fit values for the invaginated arcs in Figure 6 and Supplement figure 7.

| Sample | $R_{\max}$ [μm] | $d_m$ [μm] | $d_m / R_{\max}$ |
| --- | --- | --- | --- |
| NM IIA-KO + NMHC IIA-WT | 195 ± 16 | 110 ± 12 | 0.56 ± 0.03 |
| NM IIA-KO + NMHC IIA-ΔACD | 184 ± 31 | 134 ± 26 | 0.72 ± 0.04 |
| NM IIA-KO + NMHC IIA-ΔIQ2 | 180 ± 31 | 106 ± 23 | 0.58 ± 0.04 |
| NM IIA-KO + NMHC IIA-ΔNHT | 112 ± 28 | 44 ± 17 | 0.38 ± 0.06 |
| NM IIA-KO + NMHC IIA-3xA | 89 ± 24 | 33 ± 17 | 0.35 ± 0.08 |

## Discussion

Our work suggests that the assembly/disassembly cycle from single NM II hexamers to a minifilament involves a highly orchestrated sequence of intermediate events that are guided by isoform-specific regulatory switches. According to current knowledge, identical RLCs bind to the different NMHC II isoforms (Beach and Hammer, 2015; Billington et al., 2013; Heissler and Manstein, 2013; Sellers and Heissler, 2019), resulting in an isoform-independent activation. In such a linear system, depleting one of the paralogs would correlate with a corresponding reduction in pRLC intensity. In marked contrast, we observed a strong asymmetry between RLC phosphorylation and minifilament assembly. While all isoforms are phosphorylated *in vitro* with roughly the same efficiency (Billington et al., 2013), the spatial frequency of Ser19 pRLC *in cellula* scales linearly with NM IIA, but only to a lower degree with NM IIB and not at all with NM IIC. As the only isoform, NM IIA nearly silenced RLC phosphorylation upon its depletion. In line with this, Hu and colleagues showed that the FRAP recovery is similar for RLCs and NMHC IIA (Hu et al., 2017). Although NM IIA is the most abundantly expressed isoform (Bekker-Jensen et al., 2017; Taneja et al., 2020), these results cannot be explained by quantitative expression levels, as we have shown that overexpressing NM IIB to a level equivalent to endogenous NM IIA does not rescue this defect (Weissenbruch et al., 2021). Importantly, this does not mean that RLCs, associated with NM IIB, are not phosphorylated at all, as we show here that there is a tendency of NM IIB overexpressing cells to increase pRLC signal intensity, however, they do so to a significantly lower quality as for the reconstitution of NM IIA.

From these results, we conclude an initiating function for NM IIA, ‘upstream’ of the other paralogs. The formation of nascent minifilaments is significantly higher for NM IIA than for the other paralogs. This in turn has consequences for the assembly properties of the ‘downstream’ paralogs. Although NM IIB (and also NM IIC) minifilaments can assemble on their own, it is more likely for NM IIB hexamers to co-assemble into NM IIA pioneer minifilaments, resulting in the formation of heterotypic minifilaments (Beach et al., 2014; Shutova et al., 2014). These results are in excellent agreement with published data, which suggest that NM IIA templates for NM IIB during the assembly of heterotypic minifilaments (Fenix et al., 2016; Shutova et al., 2017; Taneja et al., 2020) and the formation of actomyosin bundles (Vicente-Manzanares et al., 2011). We conclude that the formation of heterotypic minifilaments via NM IIA is a hallmark feature in the contractile actomyosin system, even during the maturation of cardiomyocytes (Fenix et al., 2018), and thus suggest that heterotypic NM IIA/B minifilaments comprise the basic contractile unit in mammalian nonmuscle cells.

Mechanistically, the initiating function of NM IIA could be accomplished by a preferential activation and/or a higher assembly efficiency. A prior activation was previously suggested, arguing that phosphorylation of RLCs might be higher, if they are attached to NMHC IIA, rather than NMHC IIB (Sandquist et al., 2006). Although additional phosphorylation sites at Tyr155 and Thr18 inhibit RLC association to NMHC IIs (Aguilar-Cuenca et al., 2020; Gallis et al., 1983) or prolong ATPase activity (Umemoto et al., 1989), respectively, no kinases or phosphatases are known to discriminate between identical RLC molecules, when bound to different NMHC II paralogs. Of note, theoretical studies suggest that the charge pattern in the tails of NMHC IIA molecules favor their homotypic assembly over a heterotypic assembly, in contrast to NMHC IIB, for which heterotypic assembly is favored over homotypic (Kaufmann and Schwarz, 2020). This way, NM IIA and NM IIB could indeed be activated to a similar extent, but NM IIA would be more prone to assemble into homotypic minifilaments, which in turn would prevent it from RLC dissociation via Tyr155 phosphorylation (Aguilar-Cuenca et al., 2020). NM IIB would not assemble at the same activation level, but would incorporate into the nascent NM IIA minifilaments. However, we do not exclude that both effects, preferential activation and a stronger assembly of NM IIA, might act in concert.

While NM IIA dominates initial assembly, NM IIB plays a crucial role during the maturation process. It is a well-established concept that the exchange dynamics are much slower for NM IIB than for NM IIA (Sandquist and Means, 2008; Shutova et al., 2017; Vicente-Manzanares et al., 2008; Vicente-Manzanares et al., 2007). Our results show, to our knowledge for the first time in living cells, that this difference is facilitated by a load-dependency, which is encoded in NM IIB rather than NM IIA. These results are in agreement with *in vitro* experiments, showing that high load prolongs the lifetime of bound smooth muscle myosin (Veigel et al., 2003) and NM IIB (Kovacs et al., 2007), and prove a similar behavior for NM IIB but not NM IIA *in cellula*. Since blebbistatin is believed to slow down the phosphate release during the force-generating step in the crossbridge cycle, the NM II heads are stalled in the weakly bound conformation (Kovacs et al., 2004). Although NM II minifilaments do not generate force in this step, they are still attached to actin and exposed to resistive load. This load-dependency of NM IIB might fine-tune the processivity of the heterotypic minifilaments (Melli et al., 2018; Nagy et al., 2013). We hypothesize that heterotypic NM IIA/B minifilaments transition from slip to catch-bond like behaviors in regard to the ratio of active NM IIA and NM IIB in the heterotypic minifilament, engaging a clutch-like behavior (Aguilar-Cuenca et al., 2014; Case and Waterman, 2015), where NM IIB stays bound for longer periods of time, once the resisting tension reaches a certain level. It is interesting to note that this mechanism only works if a sufficient amount of actin filaments are present, which *in cellula* is typically ensured by the Rho-signaling pathway that not only activates myosin II, but also formins (Zaidel-Bar et al., 2015).

It remains to be investigated which protein domains exactly facilitate this load-dependent behavior of NM IIB. Tension above a certain threshold could lead to a conformational unfolding or stretching of NM II portions and alter the binding pockets for regulatory molecular switches, similar to proteins in FAs (del Rio et al., 2009). Several publications showed that the motor domains determine the tension contribution, while the tail domains determine the isoform-specific localization (Sandquist and Means, 2008; Taneja et al., 2020). This gives rise to possible interference points in the coiled-coil and the NHT. It was shown that phosphorylation of a serine motif, close to the NHT of NMHC IIB, promotes NM IIB assembly, while its insertion in NMHC IIA endowed NM IIA with NM IIB-like properties (Juanes-Garcia et al., 2015). However, our modeled FRAP data suggest that the observed effects might significantly rely on the crossbridge cycle rates, stored in the head regions of the NM II isoforms. In this regard, it is noteworthy that double-pRLCs (Thr18 and Ser19) are associated with long-lived NM II structures in the cell rear (Vicente-Manzanares et al., 2008) or the actin cortex (Taneja et al., 2021), and that expression of di-phosphomimetic RLC mutants specifically inhibit the exchange rate of NM IIB, but not NM IIA (Vicente-Manzanares and Horwitz, 2010). Thus, double-phosphorylated RLCs might ‘mark’ NM IIB heads, which are under mechanical load and prevent them from dissociation.

Besides heterotypic NM IIA/B minifilaments, NM IIA/C minifilaments were also reported (Beach et al., 2014). However, NM IIC does not robustly form heterotypic minifilaments in our cells and shows an unconventional association with Ser19 pRLCs, indicating that this isoform might be part of a non-canonical molecular pathway, in agreement with our earlier finding that NM IIC has a untypical role during the generation of contractile forces (Weissenbruch et al., 2021). At the current stage, we can only speculate about the unconventional/non-existent association of NM IIC and pRLC signals, or the other paralogs. *In vitro* experiments showed that NM IIC binds RLCs and ELCs, and forms bipolar minifilaments that undergo the conformational unfolding upon RLC phosphorylation (Billington et al., 2013). There are, however, significant structural differences: NM IIC minifilaments only contain 14 hexamers, have a longer bare zone, and differ in regard to the charge distribution in their NHT (Billington et al., 2013; Ronen et al., 2010; Straussman et al., 2005). These features lead to different assembly properties and a lower tendency to bind actin *in vitro* (Billington et al., 2013). Therefore, NM IIC minifilaments might only transiently associate with SFs, resulting in a low fraction of molecules, which are stably bound together with the other isoforms. Additionally, *in vitro* studies suggested that minifilaments might be composed of a mixture of phosphorylated and unphosphorylated hexamers, and that the phosphorylated fraction stabilized the unphosphorylated portion against ATP-dependent depolymerization (Kendrick-Jones et al., 1987). Although this was never shown *in cellula*, it could explain the low co-localization of NMHC IIC and pRLC signals. Future studies should dissect the relationship of RLC phosphorylation and NM IIC activation in more detail.

Altogether, our data show how NM IIA and NM IIB assemble into heterotypic minifilaments and perform together in a complementary system that generates contraction patterns, which locally self-amplify through force in a positive-feedback loop, in order to modulate the transduction of vast and noisy biomechanical cues from the extracellular environment (Graessl et al., 2017; Kamps et al., 2020). While NM IIA drives the assembly of a polarized actomyosin system by inducing initial tension and guiding NM IIB distribution in the cell body, NM IIB autonomously prolongs the lifetime of the minifilament and stabilizes the NM IIA-generated tension. This way, both motors complement each other and facilitate an optimal force output during processes like cell migration (Shutova et al., 2017; Vicente-Manzanares, 2013; Vicente-Manzanares et al., 2011), cytokinesis (Taneja et al., 2020), or the cellular steady-state morphogenesis (Weissenbruch et al., 2021). We demonstrated the consequences of an imbalance between NM IIA and NM IIB by over-assembling NM IIA hexamers in heterotypic minifilaments (Breckenridge et al., 2009; Dulyaninova et al., 2007; Dulyaninova et al., 2005; Rai et al., 2017). As it was shown that contractile pulses are an intrinsic feature of NM IIA but not NM IIB (Baird et al., 2017), over-assembly of NM IIA leads to increased tension, which overpowers the regulatory elasticity of NM IIB. *In vivo*, such contractile overshoots might cause the transduction of ‘noise’ as a stable mechanical signaling input. In this regard, NM IIB might act as a noise-filter that prolongs reciprocal signal transduction above a certain threshold. We therefore conclude that the individual ratio of both isoforms tunes the contractile output, i.e. the propagation of a mechanical signal through FAs and SFs.

## Materials and Methods

### Cell Culture

U2OS WT, COS-7, A431, A549 and HCT-116 cells were obtained from the American Type Culture Collection (Manassas, USA). U2OS NMHC II-KO cell lines were generated as described previously (Weissenbruch et al., 2021). For routine cultivation, cells were passaged every 2-3 days and maintained in DMEM (Pan-Biotech #P04-03590) supplemented with 10 % bovine growth serum (HyClone #SH3054.03) at 37°C under a humidified atmosphere containing 5 % CO_2_. For experiments, cells were plated on FN-coated coverslips or micropatterned substrates and allowed to spread for 3 h.

### Transfection and constructs

Transfections were carried out using Lipofectamine 2000 (ThermoFisher Scientific #11668027) according to manufacturer’s instructions and the cells were cultivated for 48 h before the experiment. CMV-GFP-NMHC IIA (Addgene #11347) and CMV-GFP-NMHC IIB (Addgene #11348) were gifts from Robert Adelstein (Wei and Adelstein, 2000). NMHC IIA-mApple was a gift from Jordan Beach (Loyola University, Chicago, USA). pCMV-eGFP-NMHC IIA-ΔIQ2 (Addgene #35690), pCMV-eGFP-NMHC IIA-ΔNHT (Addgene #35689) and pEGFP-NMHC IIA-3xA (Addgene #101041) were gifts from Tom Egelhoff (Breckenridge et al., 2009; Rai et al., 2017). CMV-GFP-NMHC IIA-ΔACD was produced by digesting CMV-GFP-NMHC IIA with SacII and SalI, removing the final 808 bp of the coding sequence of NMHC IIA and thereby deleting the ACD.

### Fabrication of micropatterned substrates

The master structure was produced by direct laser writing (Anscombe, 2010) and serves as a mold for the silicon stamp. The pattern of the stamp resembles a sequence of crosses with different intersections, a bar width of 5 μm and edge length of 45-65 μm. Micropatterned substrates were prepared via direct microcontact printing (Fritz and Bastmeyer, 2013). Briefly, the stamp was incubated for 10 min with a solution of 10 μg ml^−1^ FN, quickly blow dried with nitrogen, and pressed onto a coverslip. Adhesion of the coverslip to the stamp secured a proper transfer of the structures. After 10 min incubation at room temperature, coverslip and stamp were separated, and passivation was carried by backfilling the coverslip with 10 mg ml^−1^ BSA in PBS for 1 h at room temperature.

### Immunostaining

Samples were fixed for 10 min using 4 % paraformaldehyde in PBS and cells were permeabilized by washing three times for 5 min with PBS containing 0.1% Triton X-100. Following primary antibodies were used: mouse monoclonal to FN (BD Biosciences, #610078), rabbit polyclonal to NMHC IIA (BioLegend, #909801), rabbit polyclonal to NMHC IIB (BioLegend, #909901), rabbit monoclonal to NMHC IIC (CST, #8189S), mouse monoclonal to pRLC at Ser19 (CST, #3675S). All staining incubation steps were carried out in 1 % BSA in PBS. Samples were again washed and incubated with fluorescently coupled secondary antibodies and affinity probes. Secondary Alexa Fluor 488-, Alexa Fluor 647- and Cy3-labeled anti-mouse or anti-rabbit antibodies were from Jackson Immunoresearch (West Grove, USA). F-Actin was labeled using Alexa Fluor 488- or Alexa Fluor 647-coupled phalloidin (ThermoFisher Scientific #A12379 and #A22287) and the nucleus was stained with DAPI (Carl Roth #6335.1). Samples were mounted in Mowiol containing 1% N-proply gallate.

### Fluorescence imaging

Images of immunolabeled samples on cross-patterned substrates were taken on an AxioimagerZ1 microscope (Carl Zeiss, Germany). To obtain high resolution images of bipolar minifilaments and heterotypic minifilaments, the AiryScan modus of a confocal laser scanning microscope (LSM 800 AiryScan, Carl Zeiss) or a non-serial SR-SIM (Elyra PS.1, Carl Zeiss) were used. The grid for SR-SIM was rotated three times and shifted five times leading to 15 frames raw data of which a final SR-SIM image was calculated with the structured illumination package of ZEN software (Carl Zeiss, Germany). Channels were aligned by using a correction file that was generated by measuring channel misalignment of fluorescent tetraspecs (ThermoFischer, #T7280). All images were taken using a 63×, NA = 1.4 oil-immersion objective.

### Quantifications

Quantification of mean fluorescence intensities were carried out by analyzing line scans along actin stress fibers or single minifilaments in the depicted region and calculating the mean intensity. Co-localization measurements were carried out blinded by measuring the intensity of individual clusters in the single channel mode, while the other channel was switched off. To calculate the ratio of NM IIA/NM IIB in different subcellular regions, intensities for both, NM IIA and NM IIB, were summed up and the percentage of each isoform was determined. Individual numbers of analyzed cells are denoted in the respective figure captions. Quantifications of *R*(*d*)-correlations were carried out by manually fitting circles to the peripheral actin arcs of the respectively transfected cells on cross-patterned substrates. The spanning distance *d* was defined as the cell area covering the passivated substrate area. In cases, where the cell was polymerizing actin along the functionalized substrate without surpassing the complete distance to the cell edges (e.g. for NMHC IIA-ΔACD expressing NMHC IIA-KO cells), only the distance of the cell body covering the passive substrate was considered.

### FRAP experiments and analysis

GFP-NMHC IIA or GFP-NMHC IIB transfected cells were seeded on FN-coated cell culture dishes (MatTek #P35G-1.5-14-C) 3 h prior to imaging. For Blebbistatin treated conditions, 50 μM photostable Para-Aminoblebbistatin (OptoPharma Ltd., Budapest, Hungary) was added to the medium 8 h prior to imaging.

FRAP experiments were performed on an LSM 800 (Carl Zeiss, Germany) equipped with a 63× 1.4 NA oil-immersion objective and operating in the confocal mode at 37°C. During imaging, the cells were maintained in phenol red-free DMEM with HEPES and high glucose (ThermoFisher Scientific #21063029), supplemented with 10 % bovine growth serum and 1 % Pen/Strep. Images were collected at pinhole 1.0 and maximum speed using the following conditions: 10 pre-bleach frames, photobleaching of the selected region using maximum laser power and 100 iterations, post-bleach acquisition with maximum speed (300 frames for GFP-NMHC IIA and 500 frames for GFP-NMHC IIB). At maximum speed, frame rates of 2-3 fps were reached.

To correct for drift, the feature detection and matching ORB-algorithm (Rublee et al., 2011) as implemented in openCV was applied to a temporal gaussian filtered image series. In slices of 20 frames features were detected and matched. Matches were used to determine a shift per frame. This shift per frame was used to align the original videos such that the regions of interest do not move. This was implemented in custom scripts. Two square regions of interest were defined in ImageJ: The bleach spot and a reference spot with similar pre-bleach intensity. In these regions the intensity was recorded as *I*_*bleach*_, *I*_*prebleach*_, *I*_*ref*_, *I*_*ref,prebleach*_the intensity of the bleached spot after bleaching, the mean intensity before bleaching, the intensity of the reference spot after bleaching and the mean intensity before bleaching respectively. The intensity was normalized and corrected for unwanted photobleaching with

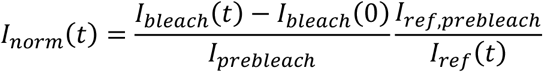

The normalized intensities were fit to *I*_*fit*_(*t*) = *δ*(1 − *exp*(−*t*/*τ*)). The fit values were reported as recovery time rand mobile fraction a.

### Modeling

FRAP trajectories of singular heterotypic minifilaments were simulated with a stochastic crossbridge and assembly model that is described in more detail in the supplemental text. We simulated for times equivalent to the experiments. The model returns trajectories of the number of fluorescent myosins in one heterotypic minifilament. Four independent trajectories were added up to obtain the FRAP intensity, which was normalized such that the initial intensity before bleaching was one. These intensity trajectories were analyzed in the same manner as the normalized experimental intensity trajectories.

## Abbreviations

ACD: assembly-competence domain
dTEM: dynamic tension-elasticity model
ELC: essential light chain
FA: focal adhesion
FRAP: fluorescence recovery after photobleaching
NHT: non-helical tail
NM II: nonmuscle myosin II
NMHC II: nonmuscle myosin II heavy chain
(p)RLC: (phosphorylated) regulatory light chain
ROCK: rho-associated kinase
SF: stress fiber
SIM: structured illumination microscopy

## Acknowledgements

This work is supported by the Deutsche Forschungsgemeinschaft (DFG, German Research Foundation) under Germany's Excellence Strategy through EXC 2082/1-390761711 (the Karlsruhe-Heidelberg 3DMM2O Excellence Cluster, to USS and MB) and EXC 2181/1 - 390900948 (the Heidelberg STRUCTURES Excellence Cluster, to USS). USS is a member of the Interdisciplinary Center for Scientific Computing (IWR) at Heidelberg. JG acknowledges support by the Research Training Group of the Landesstiftung Baden-Württemberg on Mathematical Modeling for the Quantitative Biosciences.

## Supplement figures

**Supplement figure 1:**
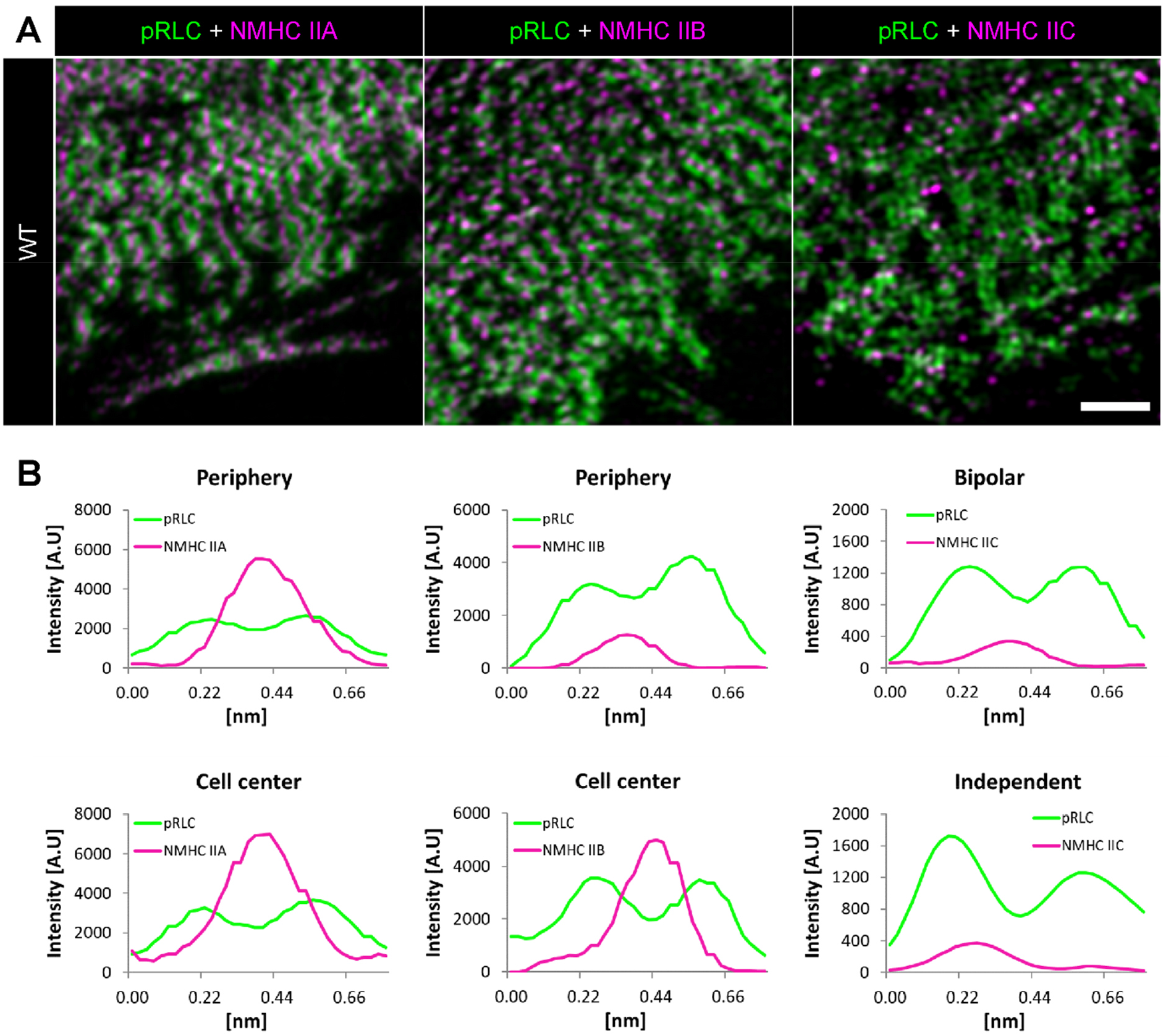
Ser19 pRLC and NMHC II paralog co-localization in bipolar minifilaments. **(A)** In WT cells, antibodies against Ser19 pRLCs label the head regions of bipolar minifilaments (green), whereas isoform-specific antibodies label the tail regions of either NM IIA, NM IIB, or NM IIC minifilaments (magenta). Fluorescence micrographs reveal a close spatial correlation for Ser19 pRLCs and NMHC IIA or NMHC IIB, while this is not the case for NMHC IIC. **(B)** Line scans from fluorescent micrographs exemplarily show the bipolar arrangement of Ser19 pRLCs and NMHC IIA or NMHC IIB signals, while Ser19 pRLCs and NMHC IIC signals do not always co-align in a bipolar fashion. Scale bar represents 2 μm.

**Supplement figure 2:**
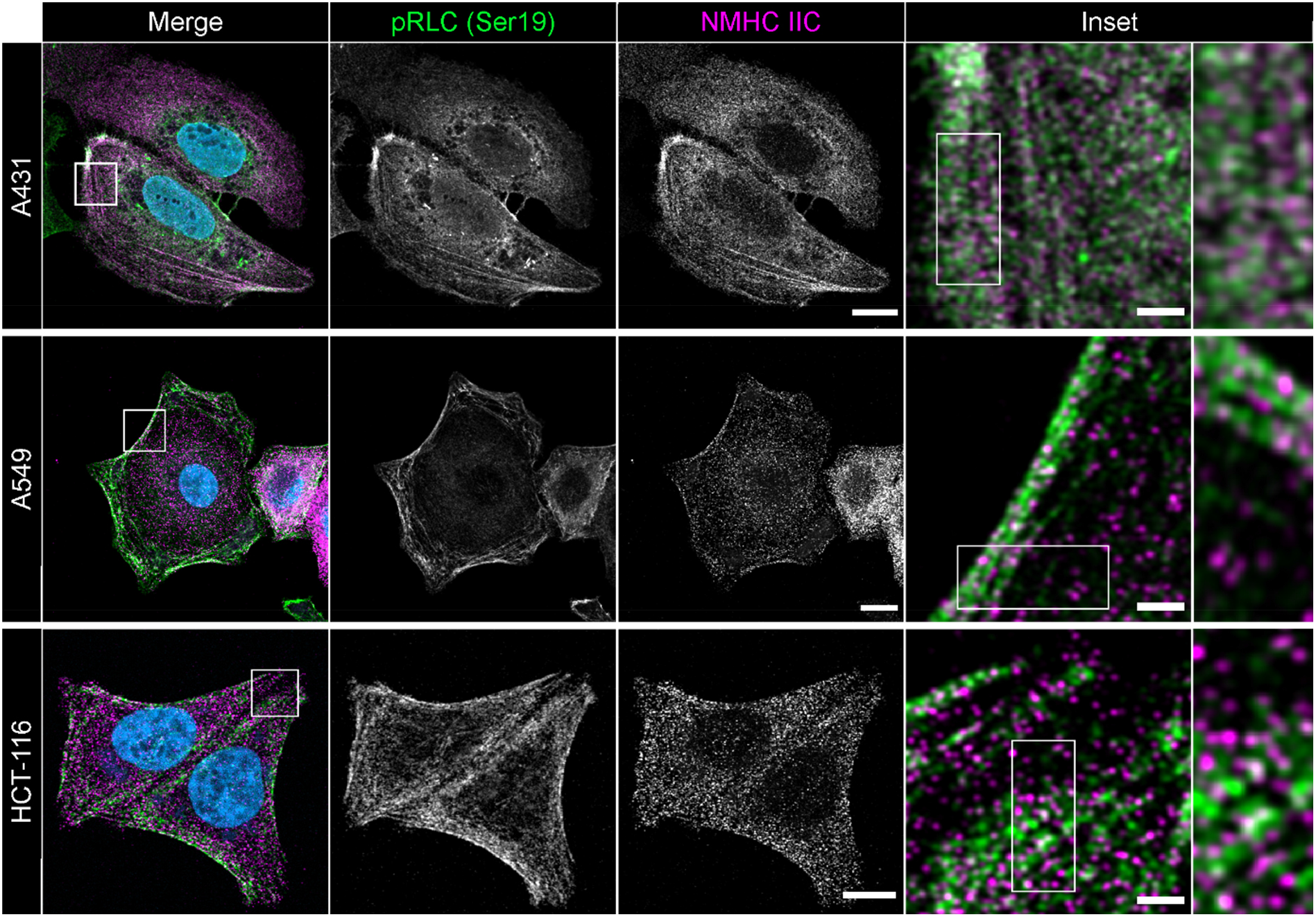
Ser19 pRLCs and NMHC IIC signals do not co-localize in a bipolar fashion. From top to bottom row: A431, A549 or HCT-116 cells were co-stained for Ser19 pRLCs (green) and NMHC IIC (magenta). Insets corresponding to the squared boxes show only few bipolar arrangements of Ser19 pRLC and NMHC IIC signals. Scale bars represent 10 μm in overviews and 1.5 μm in insets.

**Supplement figure 3:**
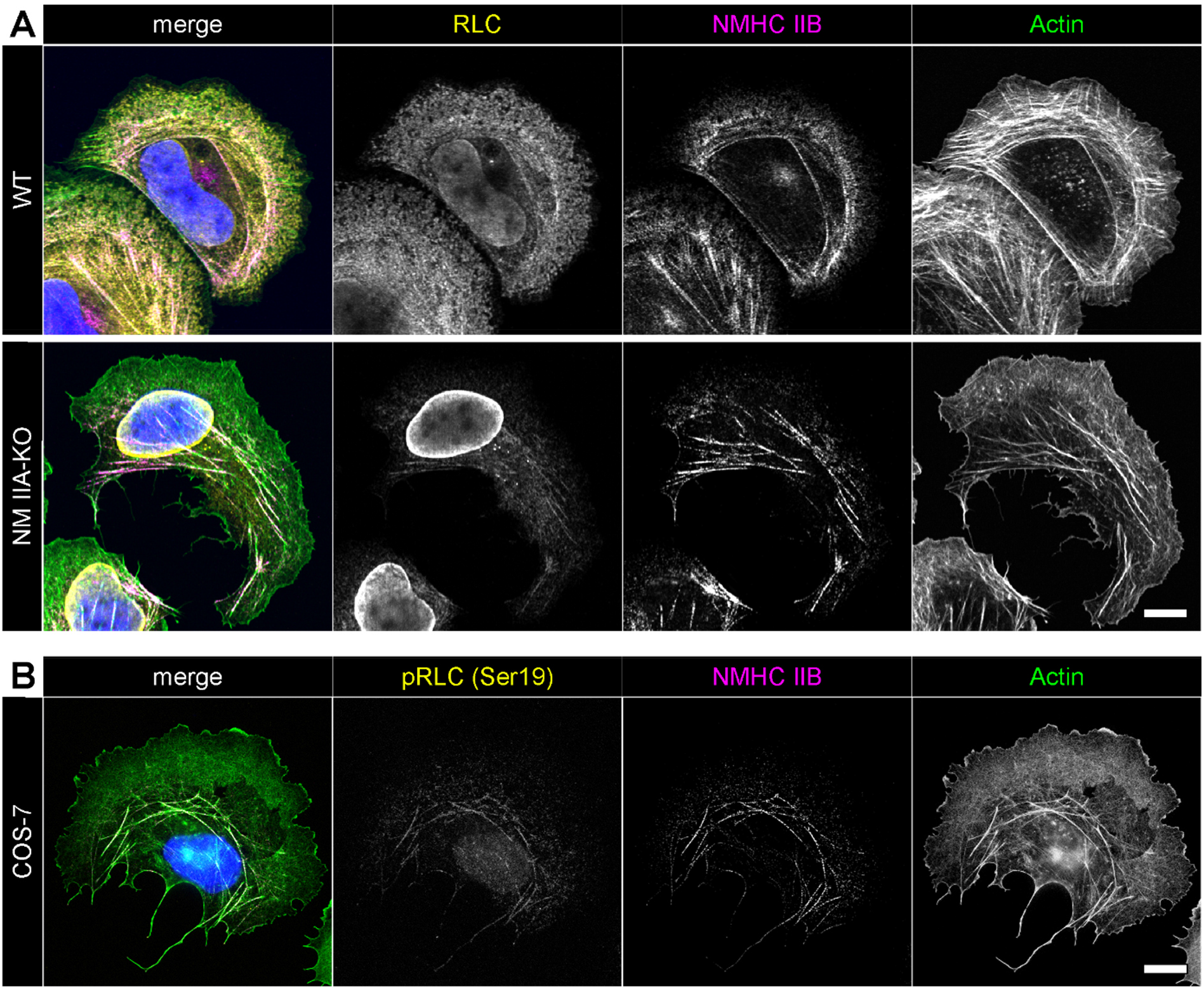
Intensity of pan RLC, Ser19 pRLC and NMHC IIB in NMHC IIA-deficient cells. **(A)** In U2OS WT cells, the pan RLC signal (yellow) is homogenously distributed and the NMHC IIB signal (magenta) is enriched in the cell center, while in U2OS NMHC IIA-KO cells, the pan RLC is almost absent and the NMHC IIB signal is densely clustered along the remaining actin fibers. **(B)** In NMHC IIA-deficient COS-7 cells, the Ser19 pRLC signal (yellow) is also almost lost and the NMHC IIB signal (magenta) is restricted to the few actin fibers. Scale bars represent 10 μm.

**Supplement figure 4:**
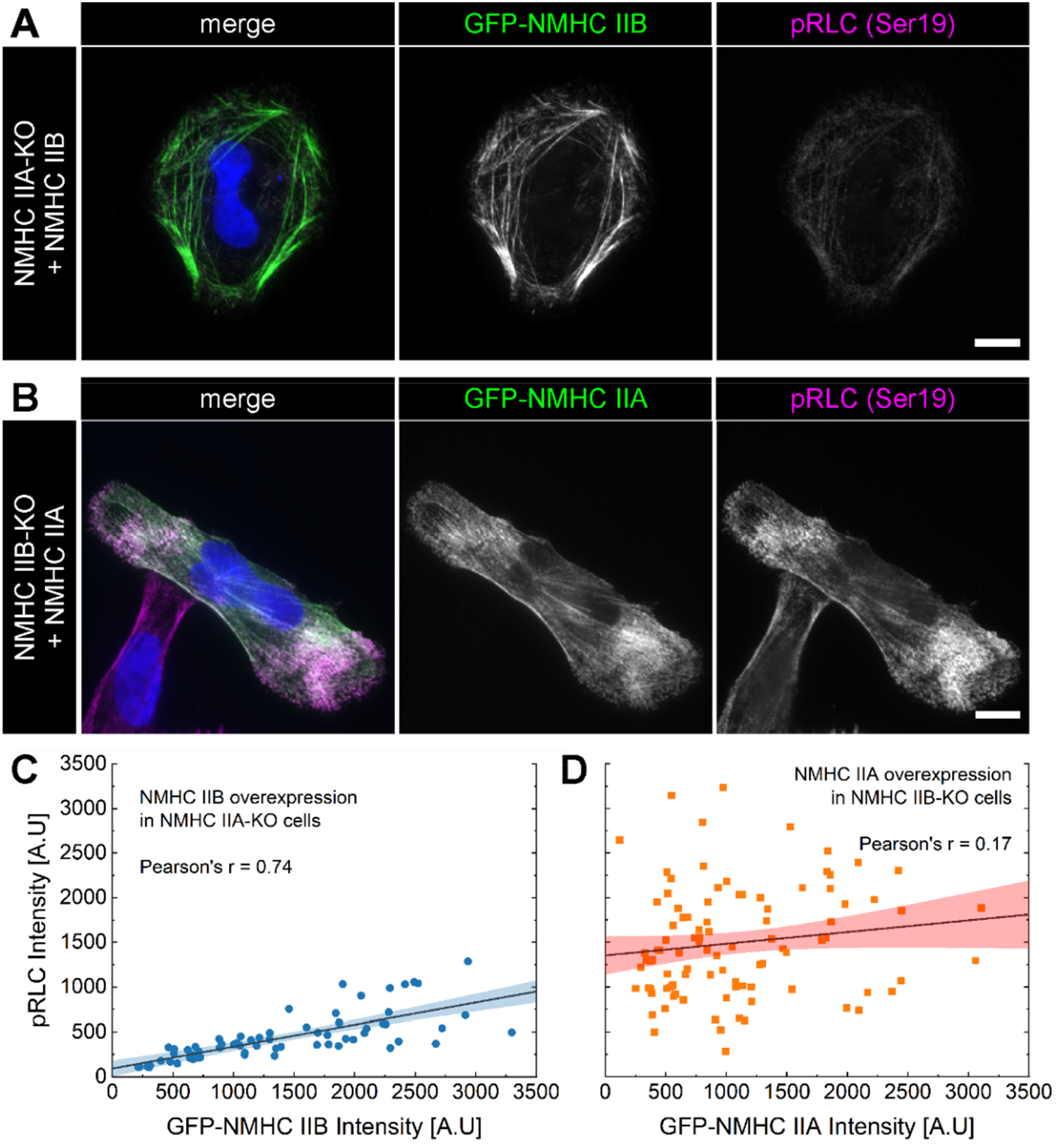
Ser19 pRLC intensity in NMHC IIA- or NMHC IIB-overexpressing cells. **(A)** Overexpressing GFP-tagged NMHC IIB in NMHC IIA-KO cells only moderately increases the Ser19 pRLC signal intensity. **(B)** Overexpressing GFP-tagged NMHC IIA in NMHC IIB-KO cells shows a strong co-localization of NMHC IIA and Ser19 pRLC signal intensities. **(C)** Plotting overexpressed GFP-NMHC IIB and Ser19 pRLC signal intensities reveals a positive correlation (pearson’s r = 0.74) with a moderate slope (data were rescaled from Figure 1E for the sake of clarity). **(D)** Plotting overexpressed GFP-NMHC IIA and Ser19 pRLC signal intensities reveals no correlation (pearson’s r = 0.17). Scale bars represent 10 μm in overview.

**Supplement figure 5:**
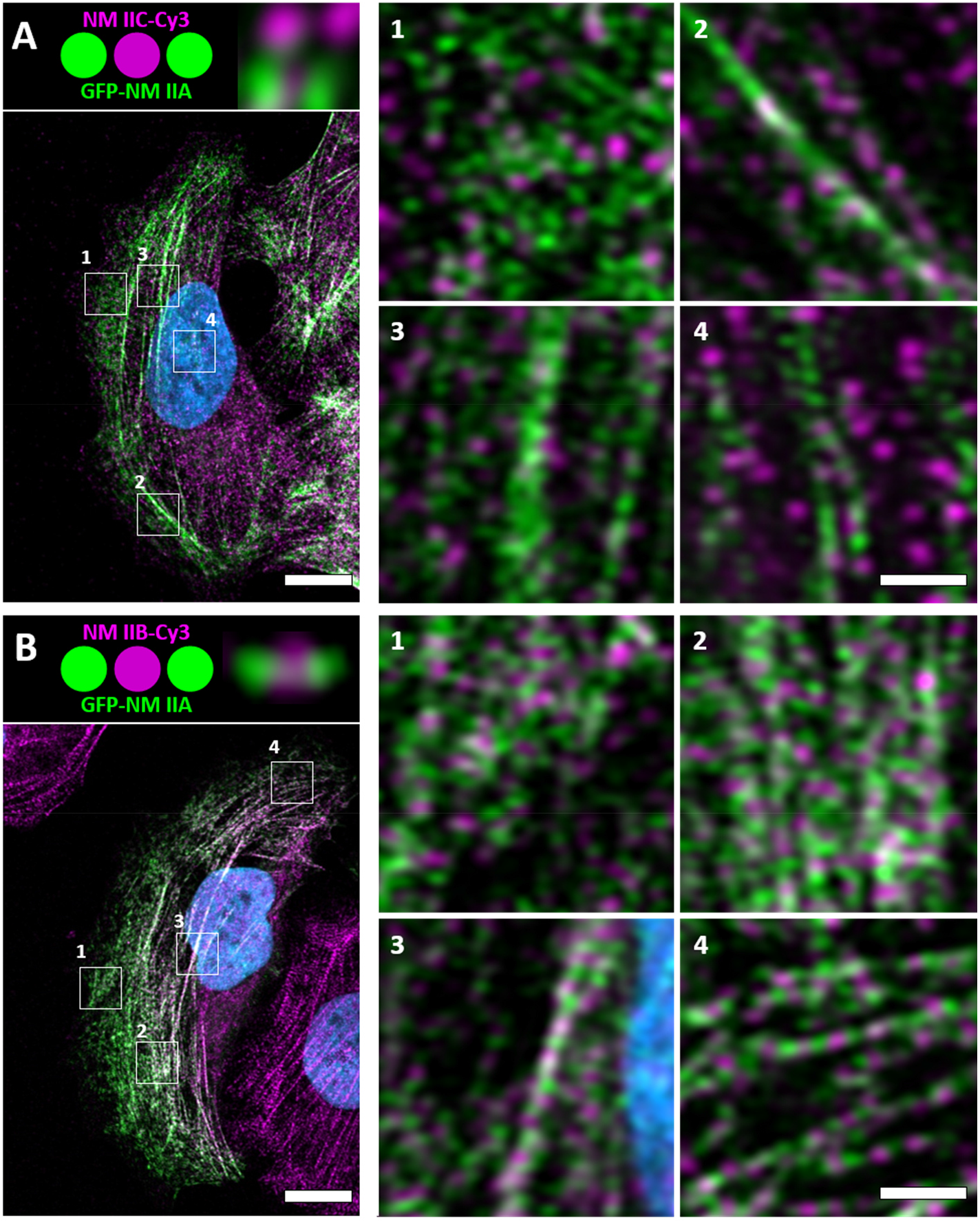
Heterotypic NM IIA/C minifilaments form less regular than heterotypic NM IIA/B minifilaments. **(A)** GFP-tagged NM IIA and endogenous NM IIC only randomly co-align in heterotypic minifilaments, independent of the subcellular position depicted in insets 1-4. **(B)** In contrast, GFP-tagged NM IIA and endogenous NM IIB robustly co-assemble in heterotypic minifilaments (compare insets 1-4). Scale bar represents 10 μm in overview images and 0.5 μm in insets.

**Supplement figure 6:**
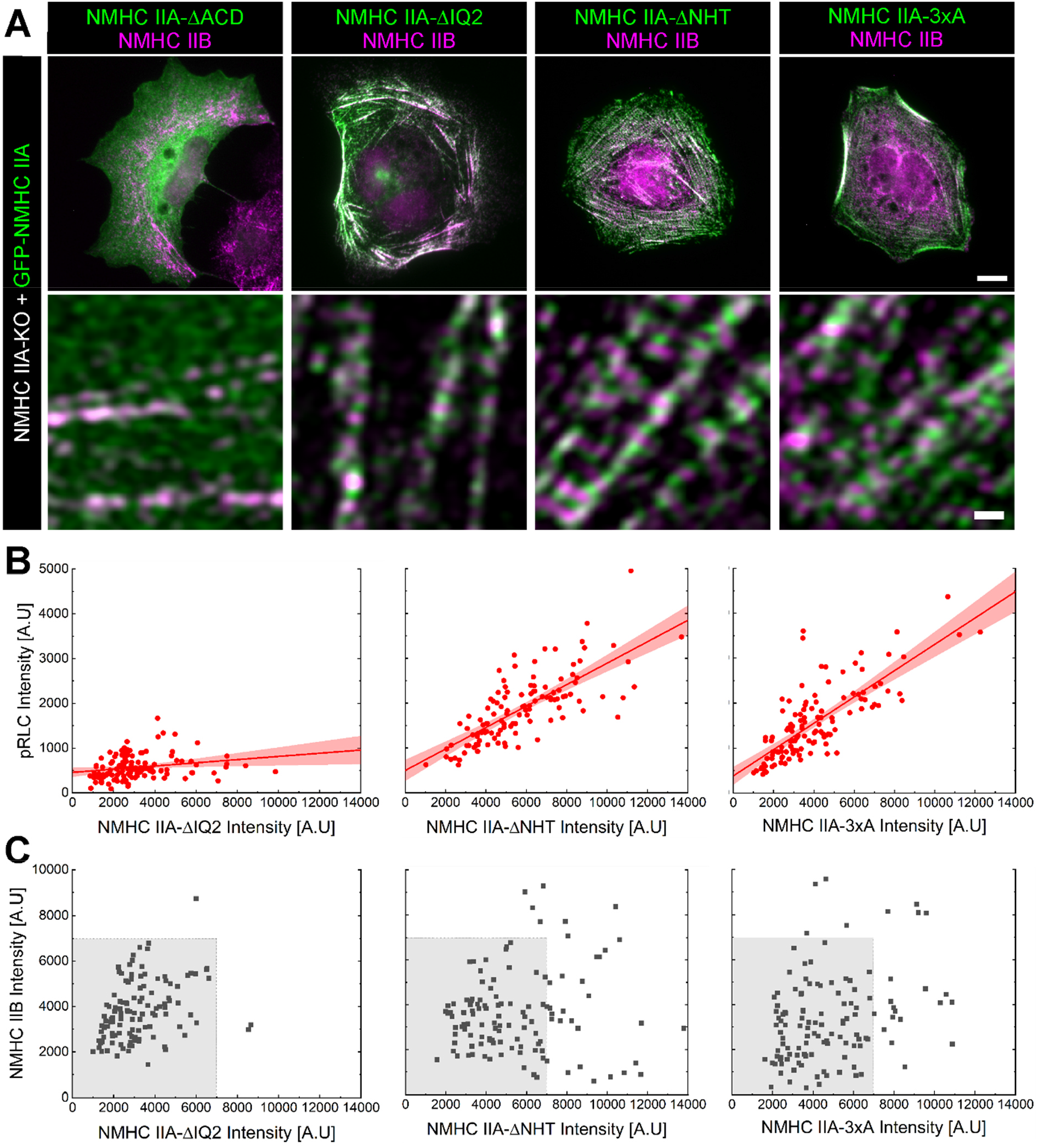
Over-assembling NMHC IIA mutants co-localize locally but not globally in heterotypic NM IIA/B minifilaments. **(A)** Expression of all NMHC IIA mutants, except for the assembly-incompetent NMHC IIA-ΔACD, induce the formation heterotypic NM IIA/B minifilaments. **(B)** Correlation of Ser19 pRLCs and the respective NMHC IIA mutants. Note that the Ser19 pRLC signal intensity does not increase in response to the expression of NMHC IIA-ΔIQ2, which lacks the binding site for the RLC. **(C)** No global correlation between NM IIA and NM IIB exists, independent of the expressed NMHC IIA construct. Note that the data in C are shifted to the lower right for NMHC IIA-ΔNHT and NMHC IIA-3xA. Scale bar represents 10 μm in overviews and 0.5 μm in details of (A).

**Supplement figure 7:**
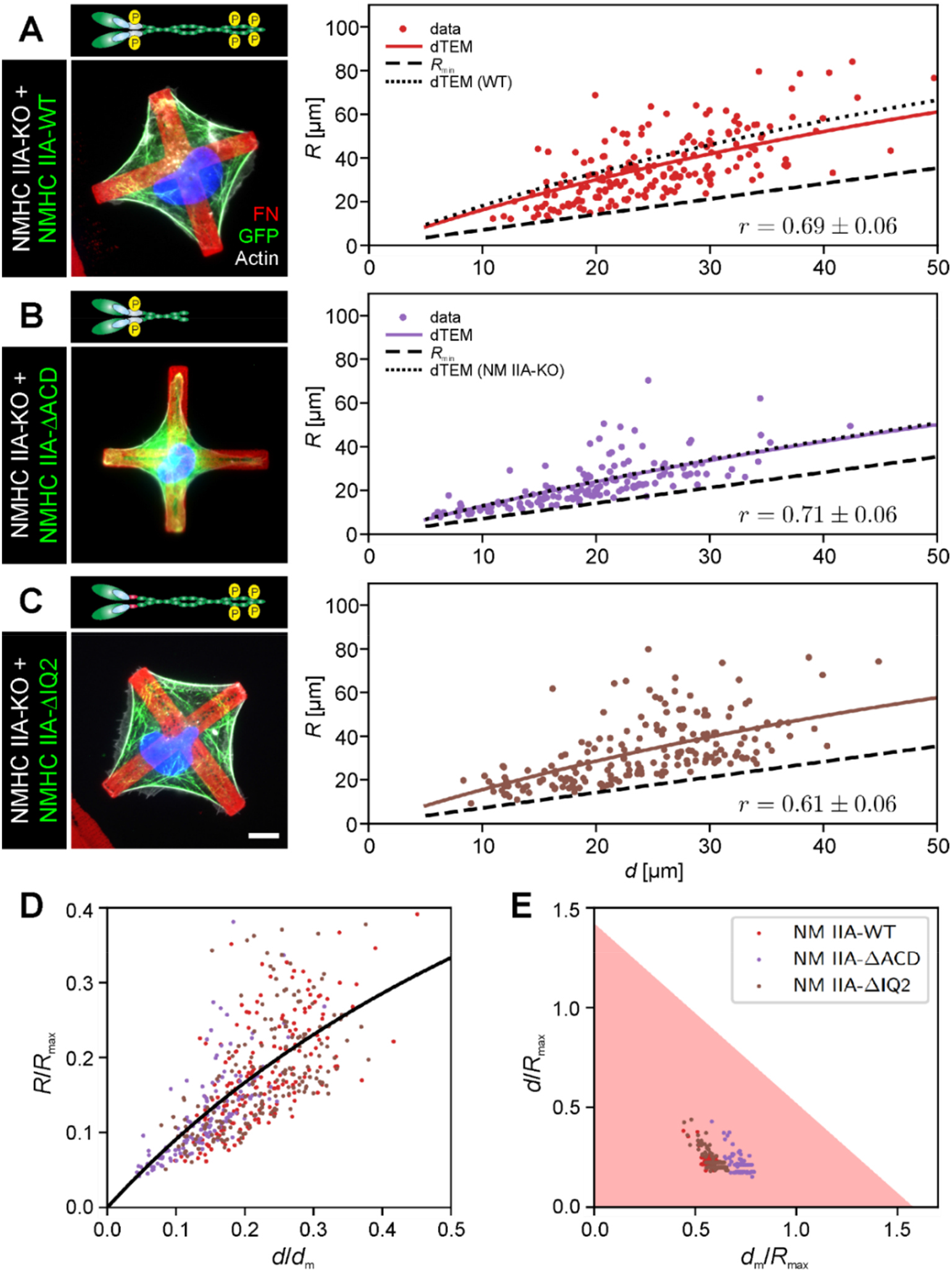
Comparison of *R*(*d*) correlation using different NMHC IIA constructs. Transfected cells were seeded on cross-shaped micropatterns to compare the *R*(*d*) correlation (correlation coefficient r given at bottom right). Solid lines denote the bootstrapped mean fit of the dynamic tension-elasticity model (dTEM), black dashed lines the geometrically possible minimal radius. **(A)** Values for NMHC IIA-WT were rescaled from Figure 6 and are shown for the sake of comparison. **(B)** Expressing NMHC IIA-ΔACD shows a comparable *R*(*d*) correlation than untransfected NMHC IIA-KO cells (compare black dashed and solid purple line). **(C)** Expressing NMHC IIA-ΔIQ2 doesn’t significantly alter the *R*(*d*) correlation compared to the WT construct. **(D)** Rescaling the experimental values with the fit parameters shows that the data of the NMHC IIA constructs lie in the linear regime of the dTEM master curve (the plateau region was cut off to visualize the overlap of the different constructs). **(E)** The ratio *d*_*m*_/*R*_*max*_ shows that NMHC IIA-ΔACD lies closest to the minimal possible radius, while NMHC IIA-WT and NMHC IIA-ΔIQ2 overlap. Quantifications were derived from N=3 experiments with n_WT_ = 54; n_ΔACD_ = 39; n_ΔIQ2_ = 50. Scale bar represents 10 μm in (A)-(C).

