## supplemental text, Supplement figure S1 for "Nonmuscle myosin IIA dynamically guides regulatory light chain phosphorylation and assembly of nonmuscle myosin IIB"

### 1 Crossbridge cycle model for nonmuscle myosin II

We model the crossbridge cycle of the different isoforms of nonmuscle myosin II with the three main mechanochemical states and the stochastic transitions between them, as depicted in Fig. S1A. Our three-state crossbridge model has been extensively tested and parametrized before [1, 2, 3, 4, 5]. Here, we extend the variant of the model introduced recently [5] to also include the effect of the myosin II inhibitor blebbistatin.

Our model works as follows. From the unbound state, myosin II heads can bind to the weakly bound state with rate $k_{01} = 0.2\,\mathrm{s}^{-1}$. Because the head is non-stereospecifically bound to actin in this situation [6], it is very prone to direct unbinding events, which occur with rate $k_{10} = 0.4\,\mathrm{s}^{-1}$. From the weakly bound state, the powerstroke occurs with the high rate $k_{12} = 1.4 \cdot 10^6\,\mathrm{s}^{-1}$. In our model, we assume this rate to be strongly reduced by the presence of blebbistatin, as it is linked to phosphate release. The effect of blebbistatin can be simulated by using $k_{12}^{\mathrm{Blebb}} = 1.5\,\mathrm{s}^{-1}$ [6]. Furthermore, the powerstroke is associated with swinging of the lever arm, which is simulated by increasing the individual motor strain $x_i$ by the powerstroke distance $d = 8\,\mathrm{nm}$. From this so-called post-powerstroke state, myosin can either return to the weakly bound state with the relatively small rate $k_{21} = 0.7$, or it can unbind from actin with a force- and isoform-dependent rate

$$k_{20}^{a/b}(F) = k_{20}^{0a/0b}[\Delta_c \exp(-k_m x_i/f_c) + (1-\Delta_c)\exp(k_m x_i/f_s)]. \tag{S1}$$

$k_{20}^{0a/0b}$ are the transition rates at zero force of the A- and B-isoform, respectively. In particular, the rate for isoform A $k_{20}^{0a} = 1.71\,\mathrm{s}^{-1}$ is much larger than the rate for isoform B $k_{20}^{0b} = 0.35\,\mathrm{s}^{-1}$, which is the only mechanochemical difference between isoform A and B in our model. The two terms in the brackets represent two different unbinding pathways, the *catch-path* and the *slip-path*. Transitions along the catch-path become slower with increasing force on the motor with force scale $f_c$. The force on the motor is calculated by the motor stiffness $k_m$ and the motor strain $x_i$. Conversely, transitions along the slip-path become quicker with increasing force with force scale $f_s$. $\Delta_c$ is the fraction of transitions following the catch-path at zero force, while the complement to one is the fraction of transitions following the slip-path. Together these two pathways constitute a catch-slip bond: At low forces the bond becomes stronger with increasing force, while at high forces the bond ruptures faster the higher the applied force. The force dependence of the rates is only taken into account if the motor is loaded against its direction of movement, the transition rate defaulting to $k_{20}^0$ for forces lower than zero.

In a minifilament, the different motor heads are mechanically coupled to form a bipolar structure with two ensembles pulling in opposite directions. We assume that in each minifilament half, $N = 15$ motor heads are active [4]. They work against external springs with strains $z_-$ and $z_+$, respectively. In addition, each side of the minifilament consists of a variable number of NM IIA and NM IIB motors, $N_a^-$, $N_b^-$, $N_a^+$ and $N_b^+$, respectively, with $N_a^- + N_b^- = N$ and $N_a^+ + N_b^+ = N$. At all times the forces acting on the myosin heads of each sides are balanced against the forces in the external springs, which yields the values for the motor strains $x_i$.

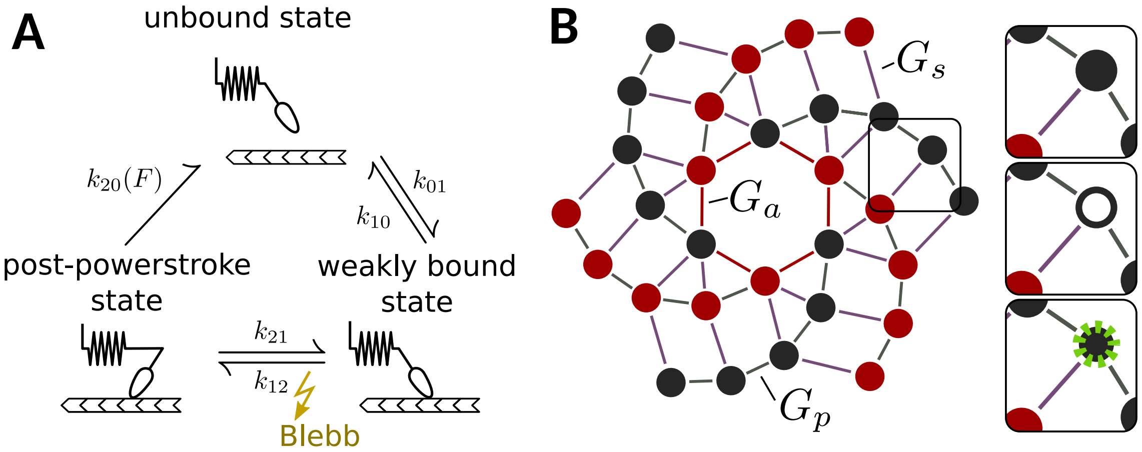


Figure S1: Model overview. (A) Mechanochemical crossbridge cycle. The rate $k_{21}$ at which the powerstroke occurs is significantly slowed down by blebbistatin, leading to a lower production of force and a lower attachment time for the individual myosin head. (B) Simulating FRAP of NM II hetero-filaments. Graph of the assembly model. Red and gray circles denote NM II hexamers with heads pointing to either direction of the minifilament, while violet, red and gray lines denote bonds with differing associated bond energies $G_a$, $G_p$ and $G_s$. The inset illustrates the two-step process of exchanging a non-fluorescent hexamer with a fluorescent hexamer (green).

### 2 FRAP simulation

In order to simulate our FRAP-experiments, we use the graph-based assembly model for myosin minifilaments introduced in [4] and extend it to include also heterotypic minifilaments. Fig. S1B visualizes the graph on which the assembly occurs. The graph follows a consensus architecture that defines the neighborhood relation of the tail domains of myosin, that are electrically charged, thereby supporting parallel and antiparallel association of two myosin hexamers to each other [7]. The assembly follows similar rules as previously introduced, i.e. the association rate of each site with neighboring occupied sites is constant and the dissociation rate of each occupied site is governed by the bindings energies of the bonds to neighboring sites that have to be broken in order to detach. The dissociation rate for one site is given by

$$k_{\text{off}} = k_{\text{off}}^0 \exp\left(-\frac{n_s G_s + n_p G_p + n_a G_a}{k_B T}\right) \tag{S2}$$

with binding energies $G_s$, $G_p$ and $G_a$, which represent the different staggers supported by the charge distribution along the myosin tail and the number of each bond type $n_s$, $n_p$ and $n_a$.

We assume that turnover is reduced by myosin heads being bound to actin [4]. In our model, each site has three possible states in total, unoccupied, occupied by NM IIA and occupied by NM IIB. The assembly model now defines separate association rates for of NM IIA and NM IIB which are defined by

$$k_{\text{on}}^{a/b} = \Delta_{a/b} k_{\text{on}}, \tag{S3}$$

where $k_{\text{on}}$ is the total association rate and $\Delta_{a/b}$ can be interpreted as the relative amounts of NM IIA and B in solution, with $\Delta_a + \Delta_b = 1$. In principle, also the dissociation rate could depend on isoform, as binding energies of the assembly depend on specifics of the charge distribution along the myosin coiled-coil [7, 8]. For simplicity, here we neglect this aspect. We use a dimensionless association-rate $\kappa = k_{\text{on}}/k_{\text{off}}^0 = 0.017$, which implies a concentration of free myosin hexamers very close to, but below the critical aggregation concentration of myosin in solution without the stabilizing effect of the crossbridge cycle [4]. This is consistent with the notion that cells only assemble myosin minifilaments if there is also actin that can be contracted, as exemplified by the architecture of the Rho-pathway. FRAP of one minifilament is simulated by associating an additional Boolean value to each occupied space indicating whether the associated myosin is fluorescent or not. The inset of Fig. S1B visualizes the two step process of replacement of a non-fluorescent myosin with a fluorescent one.

The experiments with fluorescent NM IIA and B are performed in different cell lines: NM IIA experiments are conducted with NM IIA-KO cells with reintroduced, but fluorescently tagged NM IIA, while NM IIB experiments are conducted using NM IIB-KO, with reintroduced fluorescent NM IIB. Accordingly, we consider simulations of the two systems separately. In simulations where NM IIA is assumed to be fluorescent, we assume $\Delta_a = 0.9$, while we use $\Delta_b = 0.3$ otherwise, as NM IIA is more abundantly available in the U2OS cells used here [9]. The ratio will most likely also depend on the specific cell studied. Small changes to these values however do not change the qualitative result of the simulation.

After an initial burn-in time, the number of fluorescent myosins $N^j_{\text{pre}}$ in the minifilament is recorded at one time-step before the bleach time in NM IIA and NM IIB FRAP simulations. At the time of photobleaching, all assembled myosins are set to be non-fluorescent. Newly assembling myosins of the type investigated from that point on however are fluorescent, which over time leads to an increase in the number of fluorescent myosin $N^j_{\text{fluor}}(t)$, which is also recorded. Further analysis of the simulation output is described in the material and methods section of the main text.

All model parameters are summarized in Table S1.

Table S1: Model parameters.

| Parameter | Symbol | Value | Comments |
|---|---|---|---|
| Transition rates [$s^{-1}$] | $k_{20}^{a0}$ | 1.71 | [3, 10] |
| | $k_{20}^{b0}$ | 0.35 | [3, 10] |
| | $k_{01}$ | 0.2 | [3, 10] |
| | $k_{10}$ | 0.4 | a non-stereospecific actin bound state is probed most in the presence of blebbistatin [6] |
| | $k_{12}$ | $4 \cdot 10^6$ | [3, 11] |
| | $k_{12}^{\text{Blebb}}$ | 1.5 | [6] |
| | $k_{21}$ | 0.7 | [3, 11] |
| Association rate [$s^{-1}$] | $k_{\text{on}}$ | 5 | Fit such, that NM IIA timescale matches the experiment |
| Dimensionless association rate | $\kappa = k_{\text{on}}/k^0_{\text{off}}$ | 0.017 | Below the critical aggregation without actin dynamics [4] |
| Catch-path fraction | $\Delta_c$ | 0.92 | [3, 10] |
| Isoform fractions | $\Delta_a$ | 0.9 | When simulating FRAP of NM IIA |
| | $\Delta_b$ | 0.3 | When simulating FRAP of NM IIB |
| Neck-linker stiffness [pN/nm] | $k_m$ | 0.7 | [3, 10] |
| Powerstroke distance [nm] | $d$ | 8 | [1, 2, 11] |
| Binding energies [$k_B T$] | $G_a$ | 3 | [4] |
| Binding energies [$k_B T$] | $G_p$ | 3 | [4] |
| Binding energies [$k_B T$] | $G_s$ | 1 | [4] |
| External springs [pN/nm] | $k_f$ | 4 | At high enough values, this parameter does not impact the quantitative results [12] |
